# An unusual genetic switch controls *Mycobacterium avium* pathogenesis, antibiotic resistance and colony morphology

**DOI:** 10.1101/2025.07.28.667291

**Authors:** Hannah J. Nilsson, Zoe Netter, Miles Tuncel, Nicholas E. Garelis, Nabila Siddiqui, Mira Semaan, Rita M. McCall, Kyla Dangerfield, Ella K. Cox, Laura Britt, Mark Voorhies, Allison W. Roberts, Gabriel Mitchell, Gerard A. Cangelosi, William R. Jacobs, Anita Sil, Jonathan M. Budzik, Jeffery S. Cox

## Abstract

*Mycobacterium avium* subspecies *hominissuis* (*Mah*) is an emerging environmental pathogen highly adapted to a wide range of niches, from treated water systems to mammalian tissues. On solid media, *Mah* forms two distinct colony morphologies, smooth transparent (SmT) and smooth opaque (SmO). These colony morphologies are representative of broader differential phenotypic states in which SmT cells are virulent and have high resistance to antibiotics while SmO cells are avirulent, antibiotic-sensitive and grow faster than SmT cells in culture. Importantly, *Mah* interconverts between these two morphotypes but the mechanism of SmT-SmO switching is unknown. Here we show that SmT-SmO switching is governed by a reversible transposition event that regulates expression of a periplasmic lipoprotein, Erp (<u>e</u>xtracellular <u>r</u>epetitive <u>p</u>rotein). We found that transposition of IS*1245*, an endogenous insertion sequence, into the *erp* gene correlated with the SmT-SmO transition, and its precise removal coincided with the switch back to SmT. Genetic analyses showed that *erp* is required for maintenance of the SmT state and sufficient to drive the switch from SmO to SmT. We also identified a mutation in a periplasmic protease, MarP, that locks *Mah* in the SmO state and blocks *erp*-mediated switching to SmT. Our results indicate that Erp and MarP function in a signal transduction pathway that regulates a broad transcriptional response to periplasmic stress. Moreover, identification of components that control *Mah* colony morphology switching has revealed a potential new strategy for combating the inherent antibiotic resistance of *Mycobacterium avium* infections.

## Introduction

Opportunistic pathogens adapt to extremely different habitats as they survive and grow in both environmental niches and in human tissues. *Mycobacterium avium* subspecies *hominissuis* (*Mah*) is a slow-growing bacterial pathogen closely related to *Mycobacterium tuberculosis* (*Mtb*) and is the most prevalent opportunistic mycobacterial pathogen in the United States and Europe and, alarmingly, the frequency of infections is rising^1–4^. *Mah* causes chronic pulmonary infections that are difficult to differentiate from *Mtb* infections^5^. *Mah* infections are extremely difficult to treat due to high natural resistance to antibiotics and can take years of treatment to resolve^5–7^. Unlike *Mtb*, *Mah* is found ubiquitously in the environment, especially in treated watersheds and household structures, and exist as biofilms inside water pipes and showerheads^8,9^. Although the exact route of *Mah* infection is not known, reports suggest that most infections are likely due to exposure of susceptible individuals to aerosolized contaminated water or soil^10,11^. How *Mah* adapts from an extracellular environment to become an intracellular parasitic symbiont of humans is poorly understood.

During infection *Mah* resides and grows within host macrophages and modulates cellular defenses including phagosome acidification, autophagy and programmed cell death^12–15^. These pathogenic strategies are similar to those employed by *Mtb* and enable the establishment of a replicative niche in innate immune cells^16^. As infection progresses, adaptive immunity can control bacterial replication primarily via T cell-derived interferon gamma (IFNγ), which activates macrophage defenses to restrict bacterial growth^17–19^. Loss of IFNγ results in the loss of control of both *Mah* and *Mtb*, and increases the severity of disease significantly^20–22^.

Despite the overall similarities in the intracellular pathogenic strategies and disease presentation between these related mycobacteria, there are striking differences in the molecular mechanisms that promote pathogenesis and antibiotic resistance of *Mah* and *Mtb*. For example, all of the major virulence determinants of *Mtb* known to interact with host macrophages - the ESX-1 type VII secretion system^23^, phthiocerol dimycocerosate (PDIM) and the related phenolic glycolipids^24^, sulfolipid-1 (SL-1)^25^, tuberculosis necrotizing toxin (TNT)^26^, and the secreted nucleoside TbAd^27^ - are notably absent in *Mah.* While we understand the bacterial genes necessary for *Mtb* growth in macrophages, the genes and mechanisms important for *Mah* virulence have yet to be elucidated^28^. In addition to the acquisition of virulence genes, *Mtb* has undergone other evolutionary processes that contribute to its pathogenicity that differ from *Mah*. *Mtb* has undergone genome reduction as it has adapted to surviving in a mammalian host while *Mah* has retained a larger genome, corresponding with its ability to survive in a variety of environmental niches^29^. Furthermore, the mutation frequency of *Mtb* is low and occurs primarily through the accumulation of single nucleotide polymorphisms (SNPs), likely because of strong selective pressure exerted by host defenses^30^. On the other hand, the *Mah* genome demonstrates a high degree of plasticity facilitated by large-scale genomic rearrangements, which is more characteristic of generalists that can survive in diverse environments^31,32^.

A unique aspect of *Mah* biology is its ability to switch between phenotypic states that give rise to distinct colony morphologies that correlate with virulence and antibiotic resistance^33,34^. Specifically, *Mah* can adopt three distinct morphologies: smooth transparent (SmT), smooth opaque (SmO), and rough (Rg)^33,34^. The switch to the Rg morphology is irreversible, and mediated by deletions in or near the *ser2* locus that result in loss of glycopeptidolipids on the surface of the bacteria, giving rise to cell permeability defects^35^. In contrast, switching between SmT and SmO morphotypes is reversible, indicating that it is not facilitated by permanent loss of genetic information. SmT cells demonstrate dramatically increased virulence and antibiotic resistance compared to SmO cells^36–40^. The SmT colony morphology predominates when *Mah* is initially isolated from hosts but switching to SmO occurs spontaneously as the bacteria are cultured in laboratory media. Switching from SmO to SmT is more difficult to observe during culture in laboratory media as the SmT morphology is masked by SmO cells in mixed colonies^36,37^. The reversible SmT-SmO colony morphology switch was first identified in 1965, but the mechanism that controls this switch is still unknown^41^. Understanding the genetic basis of morphotype switching is crucial for identifying bacterial factors required for *Mah* pathogenesis and antibiotic resistance.

Here we report that reversible colony morphology switching is controlled by the insertion and precise removal of the mobile genetic element IS*1245* at the *erp* locus, which encodes an enigmatic mycobacterial-specific periplasmic protein termed <u>e</u>xtracellular <u>r</u>epetitive <u>p</u>rotein (Erp). Disruption of *erp* expression alters virulence, antibiotic resistance, and the transcriptional landscape of *Mah*, correlating with changes in colony morphology. Additionally, we show that switching can only occur when the periplasmic protease MarP is functional. When MarP function is disrupted, *Mah* remains locked in the SmO morphology and is more sensitive to a broad range of antibiotics. Our results provide the first glimpse at the unique mechanism that underlies colony morphology switching and *Mah* pathogenesis and antibiotic resistance.

## Results

### Generation and characterization of smooth transparent and smooth opaque morphotypes in the *Mycobacterium avium* subspecies *hominissuis* strain mc^2^2500

To investigate the mechanism that controls colony morphology switching in *Mah*, we first derived and characterized the SmT and SmO morphologies of the clinical *Mah* strain mc^2^2500. Mc^2^2500 was isolated from an HIV-positive patient and formed SmT colonies upon initial culturing. We ensured that the bacterial populations for our experiments consisted predominantly of SmT cells by expanding individual colonies in liquid culture, creating frozen stocks, and plating aliquots on solid media to assess morphology. Only stocks that yielded greater than 99% SmT colonies were used as our starting population of SmT bacteria for the remainder of our experiments. We sequenced the mc^2^2500 genome using a combination of long- and short-read technologies and found that it consists of a 5.4 Mb genome and two plasmids, and the sequence of the *hsp65* gene confirmed that mc^2^2500 is *Mah* rather than other *M. avium* subspecies^42^. To isolate SmO cells, we passaged mc^2^2500 in liquid media and plated dilutions on solid media to identify single colonies. It was apparent that some cells had spontaneously switched from the SmT to the SmO morphology under standard laboratory culture conditions, as seen previously^33^ (Figure 1A). SmT colonies adopted both domed (Figure 1Ai) and flat (Figure 1Aii) isoforms, and their relative abundance was dependent in part on the humidity and length of incubation. Regardless of shape, every SmT colony could be identified by its translucent appearance on plates when illuminated by transmitted light. The defining characteristics of the SmO colony morphology were their solid white to yellowish color and their opaque appearance on plates when illuminated by transmitted light. SmO colonies were consistently larger than SmT colonies, and they adopted a uniform domed and rounded shape. When we visualized individual bacterial cells grown in liquid media using transmission electron microscopy, SmT and SmO morphotypes did not exhibit obvious differences from one another (Figure 1B). Both morphotypes appeared to have a similar shape and size, and there were no noticeable differences in the membrane structure between the two. There did appear to be more intracellular lipid inclusions (ILIs) in the SmO morphotype, which could indicate differences in metabolism between SmT and SmO^43^, but this was the only feature that seemed to distinguish the two morphotypes when observed at the level of individual cells. We also visualized colonies using scanning electron microcopy to see if the morphotype differences were due to multicellular organization (Figure 1C). SmT colonies formed sheet-like structures with loosely organized edges while SmO colonies appeared to be more compactly arranged with a clearly defined edge. These differences in colony organization could contribute to the macroscopic differences we observed between the two colony morphologies on a plate in which SmT colonies tend to spread out across the agar while SmO grow on top of each other. We suspect that these organizational differences result from changes in the cell surface that influence how the bacteria interact with one another.

**Figure 1.**
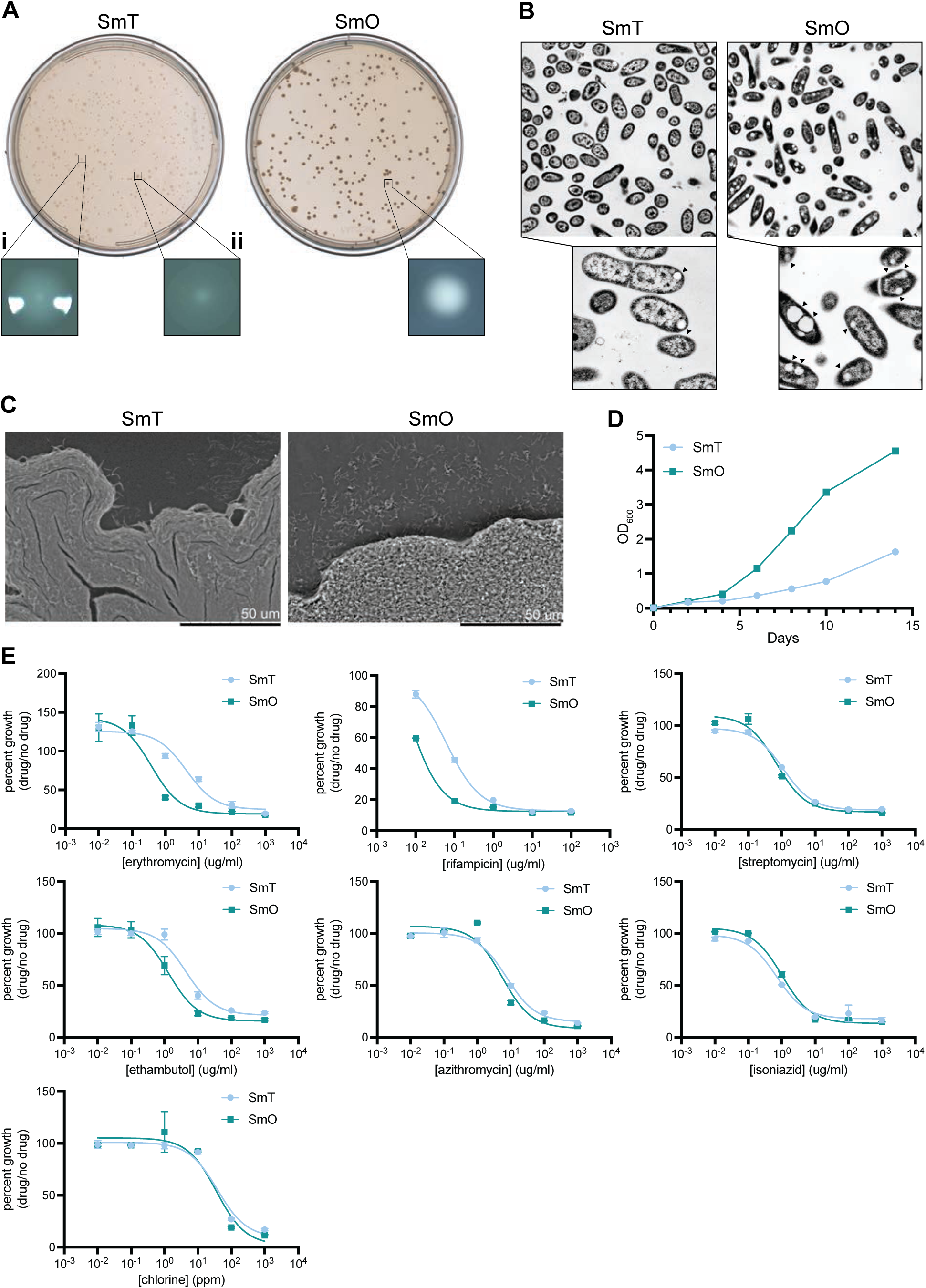
*Mah* clinical strain mc^2^2500 can adopt transparent the (SmT) and opaque (SmO) morphology. **A.** Representative images of SmT and SmO morphology for the *Mah* strain mc^2^2500 on plates captured on a scanner with transmitted light, SmT (left) and SmO (right). Single colonies of SmT (i. domed SmT and ii. flat SmT) and SmO were imaged via stereomicroscopy. **B.** Transmission electron microscopy of individual mc^2^2500 cells from SmT and SmO colonies. Arrows are used to indicate ILIs. **C.** Scanning electron microscopy of mc^2^2500 colony edges of SmT and SmO morphotypes. **D.** Growth kinetics (OD_600_ over time) of SmT (light blue) and SmO (teal) mc^2^2500 in liquid media. Each data point is the mean ± standard deviation of three technical replicates. The curve is representative of three biological replicates. **E.** Dose response curves for SmT and SmO mc^2^2500 against erythromycin, rifampicin, streptomycin, ethambutol, azithromycin, isoniazid, and chlorine, measured as OD_600_ of the culture grown with the drug at the indicated concentration divided by OD_600_ of the culture grown without the drug. Each data point is the mean ± standard deviation of two technical replicates. The curve is representative of three biological replicates.

Aside from visible changes in colony morphology, previous work showed that changes in *Mah* colony morphology are also accompanied by changes in growth, antibiotic resistance, and virulence^37–39,44^. When SmT and SmO morphotypes of mc^2^2500 were grown in liquid media, the SmO growth rate was approximately double the SmT growth rate based on OD_600_ (Figure 1D). When SmT and SmO morphotypes were exposed to antibiotics in liquid culture, SmT cells exhibited higher antibiotic resistance when compared to SmO cells to several important antibiotics (Figure 1E). The most striking difference in antibiotic resistance was made clear in the shift in the dose response curves for erythromycin and rifampicin, two first-line antibiotics against *Mah*. Specifically, the minimum inhibitory concentration (MIC) for SmT was dramatically higher than SmO for both erythromycin (64-fold) and rifampicin (16-fold) (Table 1). The difference in antibiotic resistance between SmT and SmO was more subtle in the dose response curves for streptomycin, ethambutol, and azithromycin, but the MIC values demonstrated a 2- to 8-fold increase in the antibiotic resistance for SmT in comparison to SmO for each of these antibiotics. Resistance to isoniazid and chlorine was comparable between the two morphotypes (Figure 1E, Table 1). In sum, mc^2^2500 can adopt both the SmT and the SmO morphology, and switching corresponds to specific changes in antibiotic resistance.

### Insertion and removal of the mobile genetic element IS*1245* in the *erp* genetic locus controls reversible colony morphology switching

To investigate the genetic basis for the reversible switch, we created six independent lineages of mc^2^2500 cells that had switched from SmT to SmO, and then back to SmT. We first isolated single SmT colonies to ensure that each lineage was founded by pure populations of SmT bacteria. We then isolated cells that switched to the SmO morphology by passaging SmT cells in liquid media, plating on solid media, and picking SmO colonies^36,37^. To ensure that these SmO populations no longer contained parental SmT cells, we isolated single colonies again by streaking. Previous studies have shown that the switch from SmO to SmT is rare in comparison to the switch from SmT to SmO, and we found this to be true for mc^2^2500^36,37^. Thus, to facilitate the identification of rare SmT cells that had arisen in SmO cultures, we leveraged SmT’s resistance to erythromycin to enrich for these bacteria. Specifically, we passaged our clonal populations of SmO cells in liquid culture without erythromycin to allow for spontaneous switching to occur, then added antibiotic to select against SmO cells and identified SmT colonies by plating on solid media. One full round of switching from SmT to SmO then back to SmT was considered a single lineage (Figure 2A).

**Figure 2.**
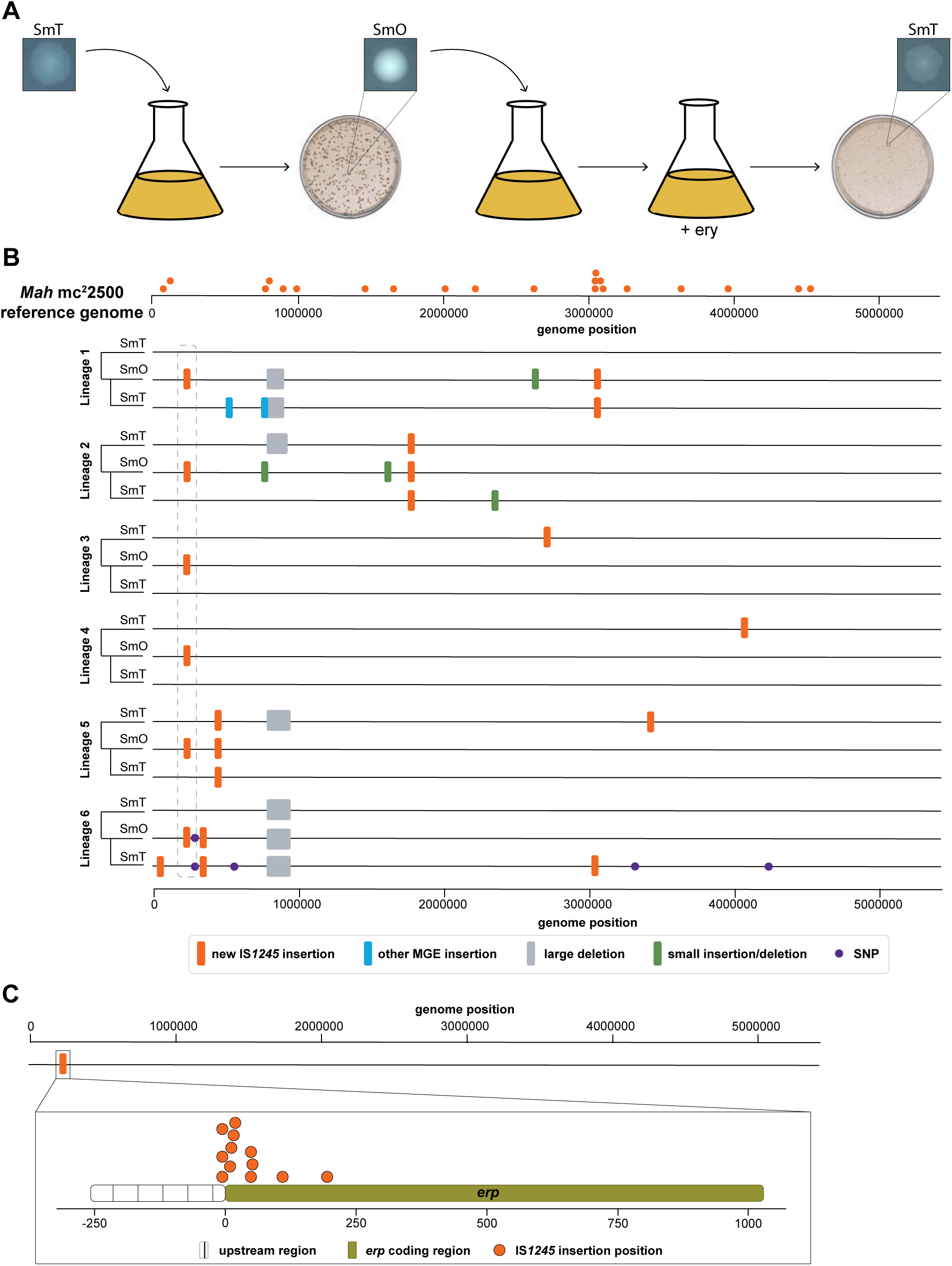
Reversible colony morphology switching depends on the insertion and removal of the transposable element IS*1245* from the *erp* locus. **A.** Schematic representation of a switching assay in which a switching lineage is generated by culturing an SmT population and plating to isolate SmO colonies that arise spontaneously. SmO colonies are cultured then treated with erythromycin (ery) before plating to identify rare switching events that revert bacteria to the SmT morphotype. A switching lineage is defined as a full round of SmT to SmO to SmT morphology switching. **B.** Whole genome sequence alignments of six independent switching lineages of mc^2^2500. Each genome was aligned to the mc^2^2500 reference genome. Orange dots above the reference genome denote IS*1245* insertions in the reference genome. New insertions of IS*1245* (orange boxes), insertions of other mobile genetic elements (blue boxes), large deletions (grey boxes), small insertions or deletions (green boxes), and small nucleotide polymorphisms (purple circles) are marked with their respective genomic position (bp) in each individual genome sequence. Dashed grey box indicates the *erp* locus in the mc^2^2500 genome. **C.** Positions of unique IS*1245* insertions in the *erp* locus derived from independent SmO colonies.

To determine if changes in the mc^2^2500 genome are responsible for the reversible switch between SmT and SmO morphologies, we isolated genomic DNA from each population of the six independent linages, performed nanopore sequencing, and compared the whole genome sequences of the SmT and SmO isolates within each lineage (Figure 2B). We identified a substantial number of transposition events and a small number of single nucleotide polymorphisms (SNPs) and insertions/deletions (indels) both within and between lineages, consistent with previous findings^31,32,45,46^. The transposable element IS*1245* contributed to discrepancies between isolate genomes in every lineage that we generated. IS*1245* is a cut-and-paste transposon that is prolific and specific to *M. avium*^47^; there are 21 intact copies of IS*1245* inserted throughout the genome of mc^2^2500. Aside from new IS*1245* insertions, another prevalent rearrangement was an approximately 100 kB deletion near position 0.8 Mb in the genome that resulted in the loss of a region containing many duplicated sequences including accessory metabolic genes, hypothetical genes, and mobile genetic elements (MGEs). In two of the four lineages where this deletion is observed, it is present in the parental SmT population but not in the subsequent SmO and SmT populations of the lineage. We hypothesize that this deletion occurred during culturing prior to isolation of genomic DNA. This example further highlights the genomic plasticity of *Mah*, which is in direct contrast to other mycobacteria like *Mtb*, whose primary source of genetic diversity is mediated by the slow accumulation of SNPs over time^48–50^.

Although each lineage exhibited varying degrees of genomic rearrangement, the only feature shared among all SmO genomes from each lineage was an insertion of IS*1245* within the genetic locus *erp* (Figure 2B, dashed box), which encodes the mycobacterial-specific <u>e</u>xtracellular <u>r</u>epetitive <u>p</u>rotein (*erp*). These independent IS*1245* insertions were absent in both the parental SmT isolates and the SmT isolates derived after the switch back from SmO, indicating that precise removal of the transposon repaired the *erp* locus. We isolated six additional SmO colonies that had similar IS*1245* transposition events in the *erp* locus. All of the IS*1245* insertions that we characterized were confined to either the upstream promoter region of the *erp* gene or the 5’ end of the open reading frame (Figure 2C). This indicated that IS*1245*-mediated disruption of *erp* expression in *Mah* is responsible for the switch from SmT to SmO, and that precise removal of IS*1245* can repair *erp* to restore the SmT phenotype.

### *Erp* expression is required for maintaining the SmT morphology in *Mah*

*Erp*, a putative periplasmic lipoprotein^51^ unique to mycobacteria, is important for virulence in pathogenic mycobacteria including *Mtb,* but its mechanism and role in *Mah* pathogenesis is unknown^52–55^ (Supplementary Figure 1). To determine if *erp* is required for the SmT phenotype, we performed complementation tests by expressing an intact *erp* gene with its native promoter on an integrating plasmid (_p_*erp*^WT^) in SmO bacteria that contained an IS*1245* insertion in the *erp* locus (*erp^−^*). These cells formed SmT colonies, indicating that Erp expression is sufficient to drive the switch from SmO to SmT (Figure 3A). Erp expression also restored antibiotic resistance, though the MIC values for *erp^−^* + _p_*erp*^WT^ consistently showed two-fold lower resistance than wild type SmT bacteria, likely due to incomplete complementation (Figure 3B, Table 1).

**Figure 3.**
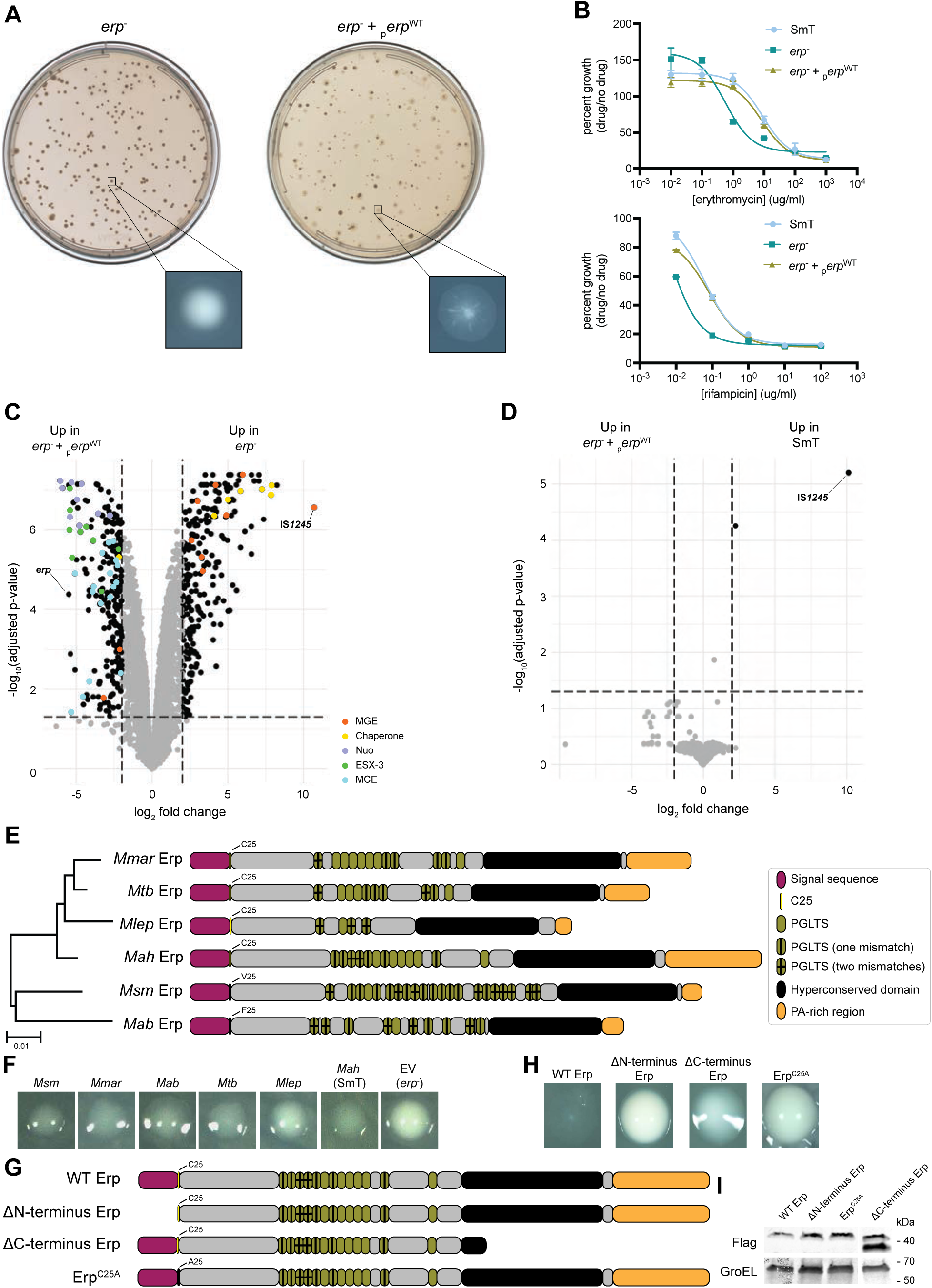
Erp is a conserved, periplasmic protein that is involved in the maintenance of the SmT morphology. **A.** Representative images of *erp*^−^ and *erp*^−^ + _p_*erp*^WT^ colony morphology generated by scanning plates with transmitted light. Single colonies of *erp*^−^ and *erp*^−^ + _p_*erp*^WT^ were imaged via stereomicroscopy. **B.** Dose response curves for SmT (light blue), *erp*^−^ (teal), or *erp*^−^ + _p_*erp*^WT^ (khaki) against erythromycin and rifampicin measured as OD_600_ of the culture grown with the drug at the indicated concentration divided by OD_600_ of the culture grown without the drug. Each data point is the mean ± standard deviation of two technical replicates. The curve is representative of three biological replicates. **C.** Volcano plot of differentially expressed genes in *erp*^−^ + _p_*erp*^WT^ compared to *erp*^−^. Adjusted p-value ≤ 0.05 and log_2_ -fold change ≥ 2.0 were considered significant and represented in black and nonsignificant results represented in grey. Select gene categories of interest are highlighted in color and indicated in the legend. **D.** Volcano plot of differentially expressed genes in *erp*^−^ + _p_*erp*^WT^ compared to SmT. Adjusted p-value ≤ 0.05 and log_2_-fold change ≥ 2.0 were considered significant and represented in black and nonsignificant results represented in grey. **E.** Schematic representation of conserved protein domains in Erps from different mycobacterial species. The putative lipidation site is denoted with the amino acid. Cladogram (left) indicates the relationship between species based on 16s sequence. **F.** Stereomicroscopy of single colonies of *erp*^−^ bacteria complemented with different mycobacterial Erps (species indicated above colony image). **G.** Schematic representation of *Mah* Erp mutations and truncations. **H.** Stereomicroscopy of single colonies of *erp*^−^ bacteria complemented with *Mah* Erp mutants. The putative lipidation site is outlined with denoted with the amino acid. **I.** Western blot of Flag-tagged mutant Erps. GroEL served as a loading control and molecular weight is indicated on the right.

To assess whether IS*1245*-mediated *erp* disruption could be observed in other *Mah* strains, we collected all available genome assemblies from *Mah* strains sequenced by Illumina or Sanger short-read sequencing technology. Short-read sequencing assembly lacks the ability to properly assemble repetitive sequences including MGEs like IS*1245*, causing contigs to break at positions where these repetitive elements are inserted. We hypothesized that if *erp* was disrupted by an MGE in other *Mah* genomes, then we would observe partial *erp* sequences at the ends of two separate contigs in genome assemblies from other *Mah* strains. After using BLAST to identify the *erp* locus in 128 *Mah* genome assemblies, we found 34 strains with the *erp* sequence split between two contigs (Supplementary Figure 2A), suggesting that these strains also experience *erp* locus disruption. Unlike our observations in mc^2^2500 where insertions were restricted to the 5’ end of the *erp* locus, other *Mah* strains contained disruptions across the entire *erp* locus. We did not find any notable correlations between the source or isolation location of a *Mah* strain and *erp* completeness (Supplementary Figure 2B-E), however it is difficult to determine if the putative *erp* disruptions existed upon isolation or arose during culture for sequencing. Beyond examples of broken *erp* sequence, we also identified two *Mah* strains with MGE sequences disrupting the *erp* locus: MAH-P-4557-08 contains an IS*1245* sequence within its *erp* locus, and the *erp* sequence of 3388 is disrupted by a different MGE (IS*1601*). Together, these analyses suggest that the phenomena of *erp* disruption by MGEs is not specific to mc^2^2500 and can be inferred from and directly observed in other *Mah* strain genomes.

### *Erp* expression alters the transcriptional landscape of *Mah*

We demonstrated that *erp* is sufficient for promoting the SmT morphology, but its exact role remains unknown. To assess *erp*’s function, we first determined global differences in gene expression between SmT, *erp^−^*, and *erp^−^* + _p_*erp*^WT^ cells using RNAseq. When we compared the gene expression profiles of *erp^−^* and *erp*^−^ + _p_*erp*^WT^ cells, over 500 transcripts were differentially expressed (Figure 3C). These results were surprising initially because Erp is a periplasmic protein that has not previously been implicated in transcriptional regulation^56^, but our data indicates that disruption of this single gene results in broad transcriptional changes. Importantly, introduction of the wild type *erp* gene restored nearly all of the differentially expressed genes to levels of SmT cells (Figure 3D). The only transcript that exceeded thresholds for significant differential expression was IS*1245*, however this phenomenon remains unclear.

Because *erp* expression results in broad transcriptional changes, we used gene ontology to predict the cellular processes and pathways associated with *erp* expression. *Erp*^−^ cells exhibited increased expression of genes involved in the cellular stress responses, including genes encoding the chaperones GroEL and DnaJ^57^, and mobile genetic element (MGE) genes^58^ (Supplementary Figure 3). This was surprising given the observation that *erp*^−^ cells grow significantly faster than SmT cells. In contrast, we found that transcripts encoding products involved in membrane-localized processes including electron transport, secretion, and membrane homeostasis were enriched in the SmT morphology. Nearly all of the genes in the *nuo* operon, which encodes the mycobacterial respiratory complex I of the electron transport chain, were upregulated in *Mah* with the SmT morphology, which was of interest as it has been implicated in virulence in both *Mtb* and *Mah*^59,60^. Additionally, transcripts in the ESX-3 operon and mammalian cell entry (MCE) protein family genes were also enriched in the SmT morphology. Both of these systems are located in the mycobacterial cell wall and have been implicated in virulence due to their importance in nutrient acquisition and utilization in the host^61–64^. Taken together, this data suggests that *erp* expression in SmT cells changes both the metabolic state and the composition of the cell wall, potentially enabling *Mah* to better respond to host immune defenses.

### Erp function depends on conserved protein domains

Erp is conserved among all mycobacterial species^54^ and consists of three main domains: an N-terminal signal sequence, a central region with a variable number of PGLTS repeat motifs, and a C-terminal domain that contains a conserved region followed by a variable PA-rich region^54,56^ (Figure 3E). To test if other mycobacterial *erp* genes could modulate colony morphology in *Mah*, we performed complementation tests by expressing *erp* homologs from *Mycobacterium smegmatis* (*Msm*), *Mycobacterium marinum* (*Mmar*), *Mycobacterium abcessus* (*Mab*), *Mtb*, and *Mycobacterium leprae* (*Mlep*) in *erp^−^* cells (Figure 3E). Expression of *Mmar* and *Mtb erp* restored the SmT morphology, while expression of *Msm*, *Mab*, and *Mlep erp* resulted in a more intermediate colony morphology (Figure 3F). We tested the antibiotic resistance of each complemented strain to rifampicin and found similar trends: *Mtb erp* expression resulted in a dose response curves that restored resistance to similar levels as *Mah erp* expression, expression of *Mmar and Msm erp* where more intermediate, and *Mab* and *Mlep erp* in *erp^−^* bacteria conferred the least amount of resistance (Supplementary Figure 4). However, strains with each of the different mycobacterial *erps* exhibited increased antibiotic resistance when compared to *erp^−^* bacteria alone, indicating that *erp* function is conserved. *Mmar* and *Mtb* Erps are most similar to *Mah* Erp, whereas *Msm*, *Mab*, *Mlep* Erps vary more in their PGLTS repeats when compared to *Mah* Erp, suggesting that a specific range of PGLTS motifs is necessary for full Erp function in *Mah*. Nevertheless, these results suggest that Erp may have a conserved function in other mycobacteria beyond the maintenance of a specific colony morphology.

To understand the functional contributions of the various Erp domains, we performed complementation tests expressing mutant variants of *Mah* Erp in which either the N- or C-terminus was truncated or the cysteine that is predicted to be lipidated was mutated (Figure 3G, Supplementary Figure 1). Expression of ΔN-terminus Erp failed to complement the SmT morphology in *erp^−^* bacteria, consistent with the idea that secretion of Erp into the periplasm in *Mah* is essential for its function (Figure 3H). The importance of Erp localization in the periplasm was further highlighted when expression of an Erp with a mutation in the cysteine that is putatively lipidated (*erp*^C25A^) failed to complement the SmT morphology in *erp^−^* bacteria. Expression of ΔC-terminus Erp also failed to complement the SmT morphology in *erp^−^* bacteria, but the function of the C-terminus remains unknown in this context. In each case, lack of complementation was not due to the absence of mutant Erp protein expression (Figure 3I), indicating that all of these domains are essential for the function of Erp in *Mah*.

### IS*1245* excision from *erp* is independent of RecA-mediated homologous recombination

Because the switch from SmO to SmT depends on the restoration of an intact *erp* locus, we sought to understand how IS*1245* excises precisely from the *erp* locus. IS*1245* insertion leads to target site duplication on each side of the transposon, resulting in an 8bp direct repeat (Supplementary Figure 5A)^47,65^. We reasoned that homologous recombination (HR) between these two sites could be catalyzed by the IS*1245* transposase or by either of two mycobacterial HR pathways, the RecA-dependent and the RecA-independent single strand annealing pathways. The large number of IS*1245* elements in the genome makes it difficult to test the transposase hypothesis, but we generated two independent Δ*recA* deletion mutants in mc^2^2500 and tested them for switching from SmO to SmT. These mutants had increased sensitivity to UV irradiation, indicating that they are indeed deficient for RecA-dependent HR (Supplementary Figure 5B). However, the Δ*recA* mutants were still able to switch from SmO to SmT, demonstrating that IS*1245* is not dependent on RecA-mediated HR for excision (Supplementary Figure 5C).

### *Erp* expression and maintenance of the SmT morphology is required for virulence

To test whether *erp* expression influences the virulence of *Mah*, we infected B6 mice intraperitoneally with *Mah* SmT, *erp^−^*, and *erp^−^* + _p_*erp*^WT^. Over the course of the eight-week infection, SmT bacteria colonized and replicated within the spleens, livers, and lungs of the mice (Figure 4A). In contrast, *erp^−^* bacteria were cleared over the course of the eight-week infection in all of these organs, confirming that SmT is more virulent, as observed previously^38^. *Erp^−^*+ _p_*erp*^WT^ bacteria exhibited similar growth kinetics to SmT bacteria over the course of the infection.

**Figure 4.**
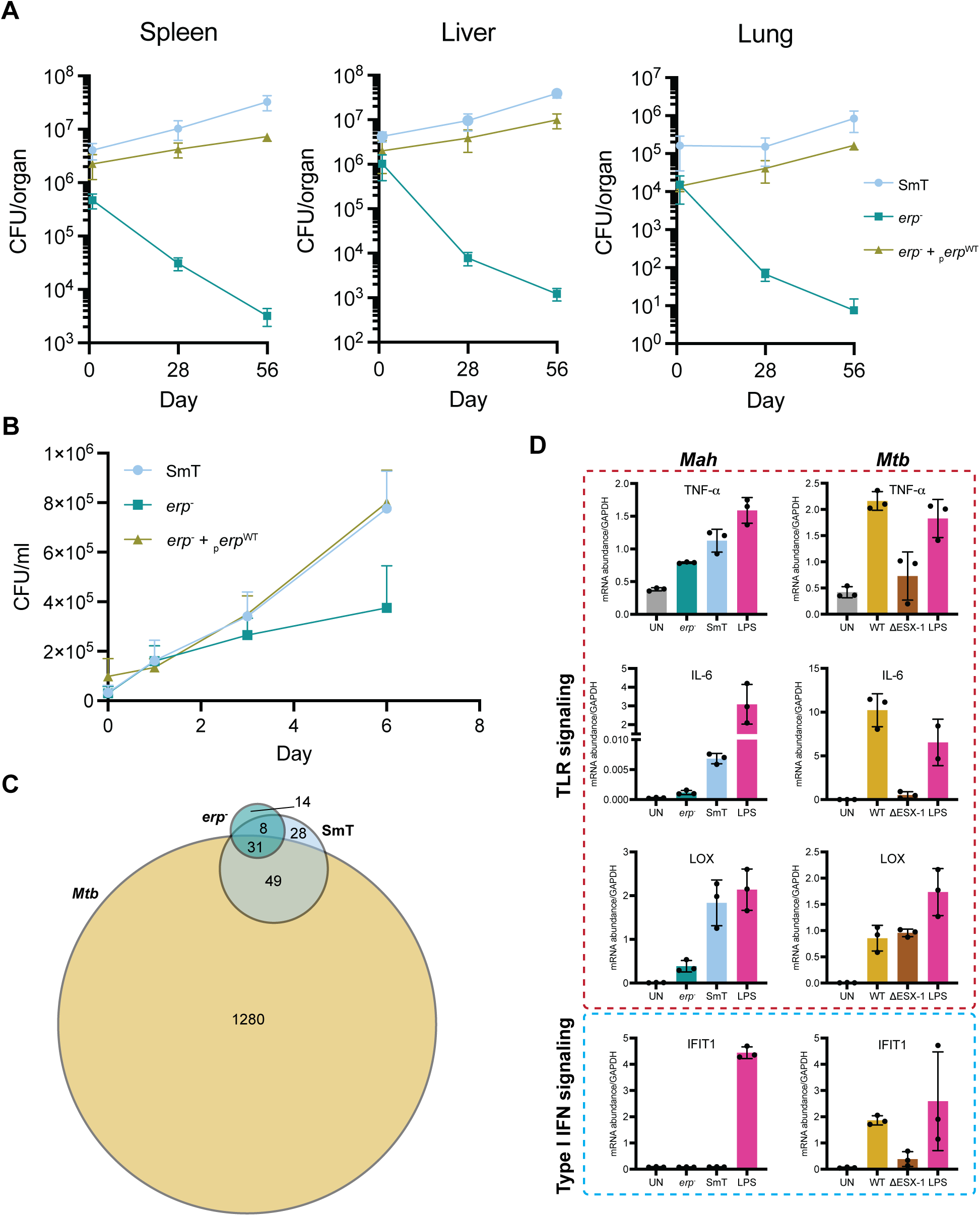
*Mah* establishes infection without triggering an overt immune response in a host in contrast to *Mtb*. **A.** Bacterial loads enumerated by colony forming units (CFU) in the spleen, liver, and lung of B6 mice infected with SmT (light blue), *erp*^−^ (teal), and *erp*^−^ + _p_*erp*^WT^ (khaki) at indicated time points. Each data point is the mean ± standard deviation of four mice per time and group. Each graph is representative of two independent experiments. **B.** Bacterial loads enumerated by CFU in murine bone marrow macrophages (BMMs) infected with SmT, *erp*^−^, and *erp*^−^ + _p_*erp*^WT^ bacteria at indicated time points. Each data point is the mean ± standard deviation of three technical replicates per time and group. The graph is representative of three independent experiments. **C.** Venn diagram comparing differentially expressed genes of SmT-, *erp*^−^-, and *Mtb*-infected BMMs 24 hours post-infection that are shared between conditions and unique to each condition. Adjusted p-value ≤ 0.05 and log_2_-fold change ≥ 1.0 (*Mah*) and ≥ 2.0 (*Mtb*) were considered significant. **D.** Reverse-transcribed quantitative PCR (RT-qPCR) of selected transcripts in SmT *Mah*, *erp*^−^ *Mah*, WT *Mtb*, and Δ ESX-1 *Mtb* infected BMMs 24 hours post-infection. mRNA expression levels are normalized to GAPDH and lipopolysaccharide (LPS) stimulation was used as a positive control. Each data point is the mean ± standard deviation of three biological replicates. Transcripts involved in toll-like receptor (TLR) signaling are outlined in red and transcripts involved in the type I interferon (IFN) response are outlined in blue.

To test if *erp* is required for growth and survival within macrophages, the primary host cell for *Mah*^66^, we infected primary murine bone marrow macrophages (BMMs) with *Mah* SmT, *erp*^−^, and *erp^−^* + _p_*erp*^WT^ bacteria. SmT bacteria grew consistently over the course of a six-day infection of BMMs while the growth of *erp^−^* bacteria was arrested after day three (Figure 4B). Importantly, *erp^−^* + _p_*erp*^WT^ bacterial growth in macrophages phenocopied growth of SmT bacteria.

To identify host macrophage responses to SmT and SmO bacteria, we infected BMMs with either SmT or *erp^−^* bacteria and harvested RNA at 24 hours post-infection, a time point where bacterial counts were equivalent, then performed RNAseq. Comparison of the transcriptional profiles between infected and uninfected BMMs identified only 108 genes that met our thresholds for being differentially expressed in SmT-infected macrophages and 52 genes that were differentially expressed during infection with *erp^−^* bacteria (Supplementary Figure 6A and 6B). The majority of host genes induced by SmO cells (38/52) were also regulated during infection with SmT cells. None of the 14 *erp^−^*-specific transcripts were associated with known immune functions and their roles in *Mah* pathogenesis remains unclear. However, the major difference between the strains was that SmT cells provoked a more robust transcriptional response than *erp^−^* cells, which likely accounts for why more genes met our significance threshold for SmT-infected cells (Supplementary Figure 6C).

The modest number of differentially expressed genes in *Mah-*infected BMMs was interesting because it directly contrasts with the broad transcriptional changes induced by *Mtb*. When comparing RNAseq data from *Mtb-*infected BMMs at the same multiplicity of infection, *Mtb* induced a 10-fold greater change in the transcriptional landscape than *Mah* at 24 hours post-infection. Only a subset of the differentially expressed genes overlapped between *Mah-* and *Mtb-*infected BMMs, but this shared subset tended to represent the largest transcriptional changes in response to *Mtb* infection (Figure 4C and Supplementary Figure 6D). This difference in transcriptional responses indicates that *Mah* and *Mtb* employ distinct strategies to achieve intracellular growth within macrophages.

To focus on innate immune gene activation, we used RT-qPCR to measure the transcript levels of genes activated by toll like receptors (TLRs) and by type I interferon (IFN), two signaling pathways activated by *Mtb*^67^. The TLR-activated genes TNF-α, IL-6, LOX, IL-1β, CXCL10, MARCO, ACOD1, and IL-1α were each induced during *Mah* infection. Transcript levels were consistently higher in SmT-infected cells compared to *erp^−^* infections, though notably less robustly than with LPS treatment (Figure 4D and Supplementary Figure 7A). This was corroborated using a cytokine bead array where we detected a small increase in TNF-α protein in the supernatants of both SmT and *erp^−^* infections (∼3-fold higher in SmT), and a very small amount of IL-6 in SmT infections only (Supplementary Figure 7B). Importantly, infection with wild-type *Mtb* elicited higher levels of all the TLR-activated genes tested than infection with *Mah* in general, indicating that *Mah* is less immunostimulatory than *Mtb*. Interestingly, wild-type and *erp^−^ Mah* did not induce expression of IFIT1, a type I interferon (IFN)-stimulated gene, which is highly responsive to *Mtb* infection. *Mah* lacks the type VII secretion system ESX-1, a known virulence factor for *Mtb* that perturbs phagosome membranes and activates cytosolic sensors to elicit a type I IFN response^68,69^. We compared type I IFN response of *Mah*-infected BMMs to that of BMMs infected with the attenuated ΔESX-1 *Mtb* mutant to assess the extent of its similarity to *Mah* infection (Figure 4D and Supplementary Figure 7A). *Mah-*infected BMMs appeared to induce even less of a type I IFN response than *Mtb* ΔESX-1, indicating that the lack of ESX-1 is responsible, in part, for the inability of *Mah* to induce type I IFN signaling in BMMs. Taken together, *Mah* is substantially less immunostimulatory than *Mtb* in BMMs, indicating that *Mah* appears to minimize early immune activation, representing a different pathogenic strategy than *Mtb*.

### Mutation in the periplasmic protease *marP* locks *Mah* in the SmO morphotype

While we were generating switching lineages, the overwhelming majority of SmO colonies were able to spontaneously switch back to the SmT morphology; however, we isolated a spontaneous mutant that remained “locked” in the SmO morphology and was no longer able to switch back to the SmT morphology (Figure 5A). By comparing the genomes of the locked SmO, SmT, and *erp^−^* bacteria, we found that the mutant had an intact *erp* locus but contained a SNP in *marP* (Figure 5B). MarP is a periplasmic protease that is involved in the response to acid stress^70–75^. Deletion of *marP* in *Mtb* results in attenuation and increased sensitivity to antibiotics^70–72^. The SNP in the locked SmO strain resulted in a missense mutation in which the glycine at position 345 is mutated to an arginine (*marP*^G345R^). The *marP*^G345R^ bacteria expressed similar levels of MarP protein as SmT and *erp^−^*bacteria, suggesting that the SNP influenced protein function (Figure 5C). Using AlphaFold to model MarP structure, we found that the G345R SNP occurs in the active site of MarP near the catalytic triad responsible for proteolysis, suggesting that this mutation could affect catalysis (Figure 5D). To test if *marP* is required for reversible switching in *Mah*, we complemented *marP*^G345R^ with a wild type copy of *marP* (*marP*^G345R^ + _p_*marP*^WT^). Unlike complementing *erp^−^* bacteria with *erp*, *marP*^G345R^ + _p_*marP*^WT^ bacteria retained the SmO morphology upon transformation with the complement construct. However, when we performed the switching assay with this complemented strain, we found that complementation restored the ability of *marP*^G345R^ bacteria to switch back to the SmT morphology (Figure 5E). Additionally, once *marP*^G345R^ + _p_*marP*^WT^ bacteria had switched back to the SmT morphology, resistance to erythromycin and rifampicin was restored to the levels of wild type SmT bacteria as indicated by the shift in the dose response curves of the complemented strain (Figure 5F). Specifically, the MIC for *marP*^G345R^ + _p_*marP*^WT^ bacteria phenocopied SmT bacteria with up to 64-fold and 16-fold higher MIC values than both *erp^−^* and *marP*^G345R^ bacteria for erythromycin and rifampicin, respectively (Table 1). Additionally, complementing *marP*^G345R^ bacteria with _p_*marP*^WT^ restored antibiotic resistance levels to streptomycin, isoniazid, ethambutol, azithromycin, and chlorine to levels comparable to wild type SmT bacteria. (Figure 1E, Table 1). Although *marP*^G345R^ is expressed in the locked SmO strain, expression of wild type *marP* restores both its ability to switch from SmO to SmT and its antibiotic resistance, suggesting that the mutation does not result in a dominant negative phenotype.

**Figure 5.**
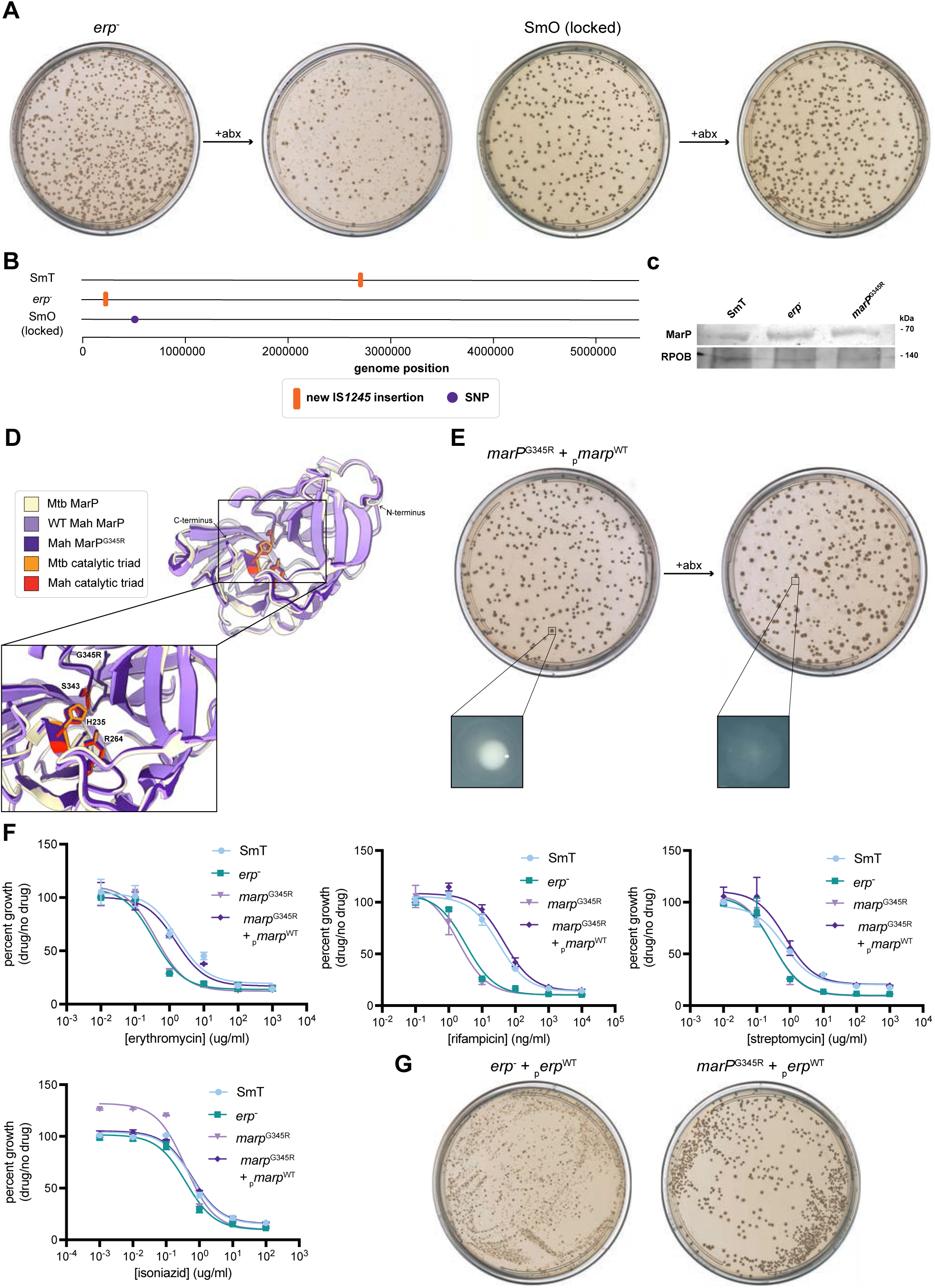
MarP is a periplasmic protease involved in the maintenance of the SmT morphology in addition to *erp*. **A.** Representative plate scans of *erp*^−^ and SmO (locked) morphology pre- and post-treatment with erythromycin. **B.** Whole genome sequence alignments of SmT, *erp*^−^, and SmO (locked). Each genome was aligned to the mc^2^2500 reference genome. Insertions of IS*1245* (orange box) and small nucleotide polymorphisms (SNPs) (purple circle) are marked with their respective genomic location in each individual genome sequence. **C.** Western blot of MarP in SmT, *erp*^−^, and *marP*^G345R^. RPOB served as a loading control and molecular weight is indicated on the right. **D.** Structure alignment of the AlphaFold model of WT *Mah* MarP (light purple) and *Mah* MarP^G345R^ (dark purple) to the crystal structure of *Mtb* MarP (tan). Insert is an enlargement of the catalytic triad. Catalytic residues are labeled as indicated in the figure legend, and G345R is labeled. **E.** Representative plate scans of *marP*^G345R^ + _p_*marP*^WT^ morphology pre- and post-switching (erythromycin treatment). Single colonies of *marP*^G345R^ + _p_*marP*^WT^ pre-erythromycin treatment and *marP*^G345R^ + _p_*marP*^WT^ post-antibiotic treatment were imaged via stereomicroscopy. **F.** Dose response curves for SmT (light blue), *erp*^−^ (teal), *marP*^G345R^ (light purple), and *marP*^G345R^ + _p_*marP*^WT^ (dark purple) against erythromycin, rifampicin, streptomycin, and isoniazid measured as OD_600_ of the culture grown with the drug at the indicated concentration divided by OD_600_ of the culture grown without the drug. Each data point is the mean ± standard deviation of two technical replicates. The curve is representative of three biological replicates. **G.** Representative plate scans of *erp*^−^ + _p_*erp*^WT^ and *marP*^G345R^ + _p_*erp*^WT^ morphology.

Because we observed a connection between different *marP* alleles and *Mah* morphotype switching capacity in the mc^2^2500 strain, we sought to define the extent of natural variation in the *marP* gene across *Mah* strains. Most *Mah* strains (∼75%) encode a MarP identical to the mc^2^2500 protein sequence, indicating that the majority of *Mah* strains should be capable of switching mediated by *erp* disruption. However, we also observed 25 strains encoding sequences that each varied by a single missense SNP, some of which were positioned in proximity to the amino acids predicted to comprise the active site (Supplementary Figure 8). Strains encoding these *marP* variants may be locked in the SmO morphology, similar to the spontaneous mc^2^2500 mutant we isolated. A small minority of strains (∼3%) contained nonsense mutations in *marP,* indicating that *marP* is not essential in *Mah*. We also observed a few *marP* sequences that, when translated, varied appreciably from the wild-type allele (∼10% amino acid identity over a portion of the C-terminal end of the protein). These variants arose from rare eight nucleotide deletions that shift the coding frame without causing early stops downstream. We hypothesize that these structural variants and *marP-*null strains would also be locked in the SmO morphotype like *marP*^G345R^, but the relative effect of active site mutations in *marP* compared to null mutants remains uncharacterized.

Both *erp* and *marP* are essential for maintaining the SmT morphotype, but they appear to function at different points in this pathway. Complementing *erp^−^* bacteria with wild type *erp* restored the SmT morphology immediately. In contrast, complementing *marP*^G345R^ bacteria with wild type *marP* did not result in immediate switching back to the SmT morphology. Instead, *marP*^G345R^ + _p_*marP*^WT^ bacteria had to undergo another round of the switching assay before the switch back to SmT occurred. This suggests that *erp* and *marP* play unique roles because *erp* expression alone can prompt the switch from SmO to SmT morphology while *marP* expression alone is not sufficient to initiate the switch.

To test whether increased *erp* expression alone could mask the SmO phenotype of *marP*^G345R^ bacteria, we performed an epistasis test and expressed *erp* in *marP*^G345R^ bacteria. Overexpressing *erp* in *marP*^G345R^ bacteria failed to restore the SmT morphology, which was in direct contrast to expressing *erp* in *erp^−^* bacteria (Figure 5G). Additionally, when we compared the gene expression profiles of *marP*^G345R^ bacteria and *marP*^G345R^ + _p_*marP*^WT^ bacteria, only 50 transcripts were differentially expressed (Supplementary Figure 9). This represents a 10-fold reduction in the number of genes that are differentially expressed between *erp^−^* and *erp^−^* + _p_*erp*^WT^ bacteria, suggesting that MarP functions downstream of the transcriptional changes controlled by Erp. This is further supported by the fact that when we compared genes that were upregulated in both *marP*^G345R^ + _p_*marP*^WT^ bacteria and *erp^−^* + _p_*erp*^WT^ bacteria, only about half the genes were shared, indicating that MarP plays its own role in this pathway in addition to the role it plays in conjunction with Erp. Taken together, Erp and MarP are essential, nonredundant components of the SmT pathway.

## Discussion

We have found that colony morphology switching in *Mah* is controlled by reversible transposon disruption of the *erp* gene. The periplasmic Erp protein is required for the SmT morphology and promotes virulence and antibiotic resistance and regulates a broad transcriptional program in the bacterium. We found that the periplasmic MarP protease is also required for these functions, and likely acts downstream of Erp, suggesting that they are components of a signal transduction pathway that controls expression of genes required for pathogenesis, antibiotic resistance and regulation of cell division (Figure 6).

**Figure 6.**
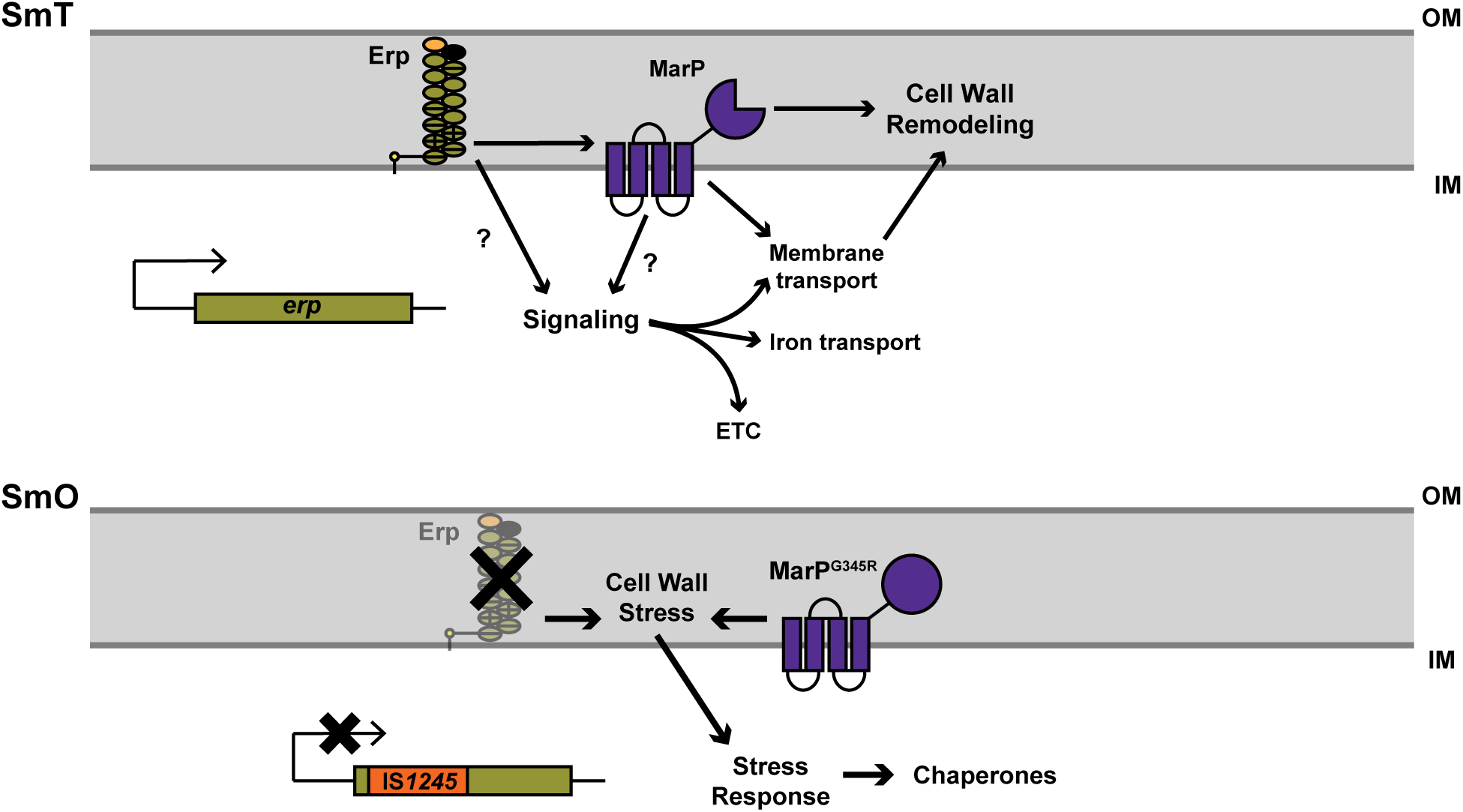
Potential mechanisms by which *erp*, IS*1245*, and *marP* mediate the switch between SmT and SmO morphologies. Top: When both *erp* and *marP* are expressed and functional, the SmT morphology is maintained. Within this context, *erp* may act in a signaling cascade, sensing external changes and inducing large scale transcriptional changes affecting metabolism, membrane transport, and other processes. Additionally, Erp and MarP may interact to promote membrane remodeling both at the transcriptional level and the membrane itself. Altogether, this could contribute to changes at the membrane that result in the SmT colony morphology and enhance *Mah*’s ability to respond to host immune system or antibiotic challenge. Bottom: When IS*1245* inserts itself into the *erp* locus or when MarP function is disrupted via mutation, the SmO colony morphology is adopted. The absence of Erp or functional MarP and resulting absence of signaling potentially leads to changes in membrane composition and cell wall stress. Cells in this state demonstrate increased growth rate and increased sensitivity to the host immune system and antibiotics. “OM”, outer membrane. “IM”, inner membrane.

Insertion and removal of the IS*1245* transposon within the *erp* locus in *Mah* may function analogously to reversible genomic alterations of other bacterial pathogens that allow for rapid adaptation to changing environments^76^. These genomic rearrangements include slipped-strand mispairing in *Haemophilus influenzae*, “gene accordions” in *Staphylococcus aureus*, gene conversion in *Neisseria gonorrhoeae*, and one other example of insertion/deletion of a transposon in controlling *Staphylococcus epidermitis* polysaccharide production^77–81^. SmT *Mah* cells have greater fitness than SmO cells during infection, but whether SmO cells have an advantage in any of its environmental niches remains unknown. While the increased growth rate of SmO cells allow them to outcompete SmT cells under ideal laboratory conditions, it seems likely that SmO cells in the wild would also be susceptible to environmental stresses, including the antibiotic-rich milieu of soil. In this case, the fitness cost of a seemingly high rate of switching from SmT to SmO may be mitigated by the elevated degree of genomic plasticity afforded by an active IS*1245* element^82^. We anticipate that our results will lead to molecular tools that will allow us to probe whether *Mah* cells in natural environmental niches exist in the SmT or SmO state.

Although *Mah* is an opportunistic pathogen and is less virulent than *Mtb,* both species share seemingly analogous pathogenic strategies – they thrive in the lungs of humans, resist macrophage defenses and grow inside these innate immune cells, are difficult to treat with antibiotics, and infections typically give rise to similar symptoms that make them difficult to diagnose accurately. However, it has become clear that the mechanistic strategies employed by these two mycobacterial pathogens are quite different. *Mtb* infections of macrophages give rise to robust TLR responses and induce the production of type I IFN via ESX-1-mediated phagosome disruption and subsequent activation of the cytosolic cGAS/STING pathway^83–85^. In contrast, we showed that *Mah* infection of macrophages gives rise to a much weaker TLR response and virtually no induction of type I IFN. The lack of a type I IFN response was perhaps expected due to the absence of ESX-1 in *Mah*, though it was possible that other mechanisms to induce the cytokine could have evolved. Interestingly, ESX-1 is required for *Mtb* to replicate in macrophages^86^, so how *Mah* replicates *in vivo* without ESX-1 remains an open question. The dampened TLR response to *Mah* indicates that it has evolved mechanisms to avoid induction of these host responses to promote its survival during infection, whereas *Mtb* can activate them without consequence^87,88^. Whether *Mah* actively inhibits TLR signaling or shields/modifies TLR ligands is unknown. Moreover, SmT cells were only slightly more immunostimulatory than SmO cells, indicating that the dampening mechanism is largely independent of Erp and MarP.

How MarP and especially Erp function in mycobacteria is not fully understood. However, several pieces of evidence suggest they are components of a periplasmic homeostatic stress response in mycobacteria and other species of the order Mycobacteriales which share the same unique cell envelope architecture. The MarP homolog in *Corynebacterium glutamicum* (42% identity with *Mah* MarP) acts as a site-1 protease and cleaves the RsdA anti-sigma factor in response to envelope stress, leading to activation of the σ^D^ regulon that serves to restore envelope integrity^89^. In *Mtb*, there are several anti-sigma factors and a known site-2 protease Rip1, but no site-1 proteases have been identified to date^90,91^. MarP regulates peptidoglycan remodeling in *Mtb* via activation of the RipA peptidoglycan hydrolase in response to acid stress, but its role in signaling in mycobacteria has not been explored^74^. Our finding that chaperone genes are activated in *erp^−^* cells and outer membrane (OM) transport complexes are downregulated is consistent with the notion that the Erp/MarP pathway in *Mah* functions to ameliorate OM stress by increasing protein folding capacity and decreasing the load of OM complexes in the periplasm. Taken together, we hypothesize that MarP in mycobacteria also acts as a site-1 protease to regulate transcription in response to envelope stress (Figure 6), though we are unable to correlate our transcriptional data with a specific stress or a known sigma factor regulon. Consistent with our genetic data, our hypothesis is that under standard conditions, Erp functions to coordinate envelope biogenesis, including assembly of multi-protein periplasmic transport complexes, with cell division. When either Erp or MarP is inactivated, uncoordinated cell division and envelope biogenesis leads to defects in transporter assembly, resulting in decreased lipid export to the OM, reduced efflux of antibiotics, and an increased growth rate.

The inherent resistance of *Mah* to many antimicrobials makes treating infections extremely difficult, resulting in multidrug treatment regimens that last more than a year^5^. Antibiotics are often poorly tolerated by patients, leading to high rates of treatment failure and relapse^92^. Our finding that MarP is required for virulence and antibiotic resistance makes it an especially attractive target for new drug development as inhibiting its function would not only increase susceptibility to host immune defenses but also increase the potency of existing antibiotics. Such inhibitors may have broad activity against mycobacterial infection, including tuberculosis and other non-tuberculosis mycobacterial infections. Moreover, there is an urgent need to better understand *Mah* pathogenesis and ecology, and to develop new strategies that will enhance current treatments for infections in humans and reduce its prevalence in natural reservoirs.

## Materials and Methods

### Bacterial strains and growth conditions

The *Mycobacterium avium* subsp. *hominissuis* (*Mah*) strain mc^2^2500 (GCA_004345205.2) was used for all experiments with *Mah*. The *Mycobacterium tuberculosis* (*Mtb*) Erdman strain was used for all experiments with *Mtb.* Liquid cultures grown in Middlebrook 7H9 media supplemented with 10% oleic albumin dextrose catalase growth supplement (OADC), 0.5% glycerol, and 0.05% Tween-80 at 37°C in a shaking incubator at 120 rpm in inkwell bottles (*Mah*) or roller bottles (*Mtb*). Bacteria were plated on solid 7H10 agar plates supplemented with 10% OADC and 0.5% glycerol and incubated at 37°C for two weeks. For growth curves, bacterial cultures in log phase growth were back diluted to an optical density (OD_600_) of 0.01 at day 0.

### Switching assay

To isolate *Mah* that switched to the SmO morphology, individual colonies of mc^2^2500 in the SmT morphology were inoculated into 7H9 media, grown to late log phase (OD_600_ 0.8-1.0), and plated on 7H10 agar. Colonies in either the SmO or the SmT morphology were struck on fresh 7H10 to isolate single colonies. To isolate colonies that switched back to an SmT morphology from the SmO morphology, individual colonies of mc^2^2500 in the SmO morphology were inoculated in 7H9 media and grown until the culture reached mid log phase (OD_600_ 0.3-0.5). These cultures were then back diluted to an OD_600_ of 0.05 in 7H9 media containing erythromycin (8 µg/ml) and grown to mid-log (OD_600_ 0.3-0.5). The culture was then plated on 7H10 and allowed to grow at 37°C for two weeks.

### Plasmid construction and strain generation

For construction of complementation plasmids, *erp, marP,* and *recA* were PCR amplified with PrimeStar GXL polymerase (Takara) from the genomic DNA of mc^2^2500. *Msm erp, Mmar erp, Mab erp,* and *Mtb erp* were amplified by standard PCR from the genomic DNA of *M. smegmatis* (mc^2^155), *M. marinum* (ATCC 927), *M. abcessus* (ATCC 19977), and *M. tuberculosis* (Erdman (ATCC 35801), respectively, including native upstream and downstream regulatory elements. *Mlep erp* (AL450380.1) was ordered as a synthetic gene block from Integrated DNA Technologies. Linear DNA, mycobacterial integrating shuttle vector pMV306, or the episomal shuttle vector pMV261^93^ were each digested with the restriction enzymes ClaI and HpaI and ligated together using T4 DNA ligase. Plasmids were transformed into and amplified with DH5α *Escherichia coli* and verified by whole-plasmid nanopore sequencing. ΔN-terminus *erp,* erp^C25A^, and ΔC-terminus *erp* were generated by utilizing the Q5 site directed mutagenesis kit (New England Biolabs) on the pMV306-*erp* vector. Electrocompetent *Mah* bacteria were prepared and transformed via electroporation as previously described^94^. Bacteria transformed with modified pMV306 plasmids were plated on 7H10 plates with hygromycin (100 μg/ml). Δ*recA Mah* were generated as previously described^95^. Transformants were plated on 7H10 plates with kanamycin (50μg/ml) and CRISPR activity was induced by plating transformed bacteria on 7H10 plates with anhydrous tetracycline (500ng/ml).

### Colony morphology imaging

Plates were imaged on an Epson Perfection V850 Pro photo scanner. Individual colonies were imaged using a Zeiss Axio Zoom V16 stereomicroscope fitted with a PlanNEOFLUAR Z 1.0x objective lens. Images were captured using an AxioCam506 color camera.

### Transmission electron microscopy

Bacteria were grown to mid log phase in liquid media and pelleted by centrifugation. Bacteria were then prepared as previously described^96^. Electron microscopy analysis was performed with the assistance of the Analytical Imaging Facility at the Albert Einstein College of Medicine of Yeshiva University on a JEOL 1200EX transmission electron microscope at 80 kV.

### Scanning electron microscopy

MCE Membrane Filters, 0.45 μm pore size (Millipore Sigma) were first sterilized in 70% ethanol and then placed on 7H10 plates. Bacteria were allowed to grow on the filters for two weeks at 37°C. Samples were then prepared as previously described^97^ and imaged with a Hitachi TM-4000 Scanning Electron Microscope.

### Dose response to antimicrobials

*Mah* was grown to log phase (OD_600_ 0.2-1.0), back diluted to an OD_600_ of 0.05, then treated with or without a ten-fold dilution series of each antimicrobial compound to a final volume of 100µl per well in 96 well, non-TC-treated plates. Plates were incubated at 37°C for six days without shaking in a moist Tupperware container to prevent evaporation. After six days, bacteria were fixed with an equal volume of 10% formalin. Optical densities were read at 600nm with a SpectraMax M3 Microplate Reader. Percent growth was calculated relative to bacteria grown in media alone for each strain. Data were fit using a dose-response inhibition fit on GraphPad Prism.

To calculate the minimum inhibitory concentration (MIC), *Mah* were grown and exposed to antibiotics in a two-fold dilution series as described above. The MIC was determined to be the concentration of antimicrobial compound at which the absorption at 600nm was at 10% as compared to the absorption at the lowest concentration of the antimicrobial compound.

### Whole genome sequencing and analysis

#### Genomic DNA extraction

*Mah* strains were grown to mid- to late-log phase (OD_600_ 0.5-1.0) in 15ml of 7H9 media. Bacteria were harvested by centrifugation, resuspended in 450µl of resuspension buffer (25mM Tris-HCL, pH 7.9, 10mM EDTA, 50mM glucose) with 50µl lysozyme (10mg/ml), and incubated at 37°C overnight. The following day, 25µl of proteinase K (20mg/ml) and 100µl of sodium dodecyl sulfate (10% w/v) were added and the suspension was incubated at 55°C for 30 minutes. A solution of 200µl of 5M NaCl and 160ul of cetrimide saline solution (0.7M NaCl, 10% cetrimide (Sigma) w/v) was added to the samples and the suspension was incubated at 65°C for 10 minutes. Genomic DNA was extracted by adding an equal volume of chloroform:isoamyl alcohol (24:1) to the samples and centrifuging the suspension for 10 minutes. The aqueous layer was isolated, 0.7 times the volume of isopropanol was added to precipitate the DNA, and the DNA was pelleted. The DNA pellet was then washed with 70% ethanol, dried, and resuspended in water.

### DNA sequencing

#### DNA sequencing and assembly

Genomic libraries of the mc^2^2500 reference strain were prepared by QB3 Genomics at the University of California, Berkeley for both PacBio and Illumina sequencing. To assemble the reference genome for mc^2^2500, PacBio and Illumina sequences were compiled and assembled using Canu. Small corrections to the assembly were made via Pilon, and the final assembly was compared to the original reads to ensure accuracy. Subsequent sequencing of mc^2^2500 derivatives for morphology lineage tracing was conducted with Oxford Nanopore technology with custom analysis and annotation performed by Plasmidsaurus.

#### DNA sequence analysis

Genomes were prepared for alignment using CLC Genomics Workbench to standardize the orientation and start point to match the mc^2^2500 reference genome. Pairwise alignments and analysis of SNPs and gaps were generated using Mauve^98^. For lineage tracing experiments, the parental SmT genome for each lineage was first aligned to the mc^2^2500 reference genome, then aligned pairwise to each of the SmO and SmT genomes derived from the parent. SNPs and gaps calculated by Mauve were validated by hand and plotted using a custom Excel macro.

### RNA sequencing

#### RNA isolation (Mah)

Individual colonies of *Mah* were grown to mid-log phase (OD_600_ 0.3-0.5) in 25ml of 7H9 media. Bacteria were harvested by centrifugation, washed once with Gibco’s phosphate buffered saline (PBS), and resuspended in 1ml Trizol before bead beating with the Savant FastPrep FP120 Cell Disruptor three times for 30 seconds at 6.5m/s. 200µl of chloroform was added to the lysate. Samples were then mixed and centrifuged for 15 minutes at 4°C. The aqueous layer was isolated, an equal volume of 70% ethanol was added, and RNA was extracted using silica spin columns (Invitrogen PureLink RNA kit). Purified RNA was treated with DNase I (10U) on column at room temperature for 15 minutes (Qiagen).

#### RNA isolation (Macrophages)

Macrophage monolayers were harvested for RNA isolation by aspirating media from the wells, washing once with PBS, then adding 1ml Trizol to each well. Trizol was pipetted thoroughly over the monolayer to dislodge and resuspend the cells. Macrophage Trizol samples were treated as described for *Mah* samples starting at the chloroform addition step.

#### RNA sequencing and analysis (Mah)

RNAseq libraries were prepared from biological triplicate samples and sequenced with Illumina HiSeq (2×150bp) by Azenta Life Sciences. Reads were mapped to the GenBank assembly for the *Mah* genome mc^2^2500 (ASM434520v2) and quantified using Kallisto pseudoaligner (version 0.44.0). The R package edgeR (version 4.0.2) was used to determine differentially expressed genes. Normalization was performed using trimmed mean of M-values. The cutoff for significantly differentially expressed genes was a log_2_(fold-change) ≥2 or ≤-2 and adjusted p-value <0.05.

#### RNA sequencing and analysis (Macrophages)

RNAseq libraries were prepared from triplicate wells using PolyA Enrichment, sequenced at a depth of 50M reads with Illumina HiSeq, and aligned to the GRC38/mm10 reference by SeqCenter. Trimming, QC, and differential gene expression analysis were performed using SarTools^99^ in Galaxy Pasteur^100^. The cutoff for significantly differentially expressed genes was a log2(fold-change) ≥1 or ≤-1 (*Mah*), ≥2 or ≤-2 (*Mtb*) and adjusted p-value <0.05. Volcano plots and heatmaps were generated with Prism.

### Bioinformatic analysis

*Mycobacterium avium* ssp*. hominissuis* strain genome sequences and isolate information were downloaded from NCBI, including all complete genomes deposited before fall of 2024. Local BLAST was performed to identify and catalog *erp* and *marP* loci in each strain. Broken *erp* sequences were identified by BLAST hits returning partial sequence homology split between two contigs. The *erp* phylogeny was generated by manually repairing broken *erp* DNA sequences, then conducting a multiple sequence alignment with MUSCLE. The resulting alignment was uploaded to MEGA to generate a minimum evolution phylogenetic tree, and a custom R script was used to generate a graphical representation of the isolate information. The mycobacterial phylogeny in Figure 3E was generated in MEGA as described above using representative 16s sequences from each species. For *marP* sequence analysis, the *marP* coding sequences were translated into amino acid sequence and aligned using Clustal Omega. This alignment was used to cluster identical *marP* alleles and create a representative set of unique MarP sequences. Next, each individual representative alternative MarP allele was aligned pairwise to the wild type sequence using Needle to validate the position and extent of mutation. Graphic representation of different MarP alleles was generated by hand.

### UV Assay

*Mah* strains were grown to early log phase (OD_600_ 0.3-0.4) in 15ml of 7H9 media. A ten-fold dilution series was performed for each strain, and dilutions were spotted onto 7H10 plates using a frogger. Plates were allowed to dry for an hour and then plates were exposed to either 0, 4, or 8mJ/cm^2^ in a Stratalinker UV Crosslinker. Bacteria were grown for two weeks at 37°C prior to imaging on a photo scanner.

### Western blot

*Mah* was grown to late-log phase (OD_600_ 0.6-0.8) in 20ml of 7H9 media in inkwell bottles. Bacteria were harvested by centrifugation at 2850*g*, washed once with PBS, resuspended in lysis buffer (1mM EDTA, 1mM PMSF, and cOmplete-mini, EDTA free (Roche) in PBS) and boiled for 10 minutes before bead beating with the Savant FastPrep FP120 Cell Disruptor three times for 30 seconds at 6.5m/s. Protein lysates were separated by SDS–polyacrylamide gel electrophoresis (Bio-Rad Miniprotean TGX 4–20%) and transferred onto nitrocellulose membranes using a semi-dry transfer method with BioRad Trans-Blot Turbo Transfer System. After blocking with Odyssey blocking buffer for 1 hour at room temperature, *erp* expression was probed using mouse anti-DYKDDDDK (Invitrogen, dilution 1:1000) and *marP* expression was probed using mouse anti-*marP* (1:2000, gift from Sabine Ehrt). Anti-GroEL rabbit polyclonal (Enzo ADI-SPS-875, dilution 1:1000) or anti-RpoB rabbit monoclonal (Abcam ab191598) were used as a loading control. Primary antibodies were incubated overnight at 4°C. Blots were washed three times with PBS and then incubated for 30 minutes at room temperature with secondary antibodies. Licor IRDye 800CW goat anti-mouse IgG (1:10000 dilution) and Licor IRDye 800CW goat anti-rabbit IgG (1:10000 dilution) were used as secondary antibodies. All antibodies were diluted in Odyssey blocking buffer. Membranes were imaged on an Odyssey scanner (Li-cor).

### Mouse infection with *Mah*

All animal studies have been approved by the University of California Berkeley Institutional Animal Care and Use Committee (protocol no. AUP-2015-11-8096-1). C57BL/6 mice were obtained from the Jackson Laboratory, Bar Harbor, ME. Female mice were infected at 6-8 weeks of age with 5.0 × 10^7^ colony forming units (CFUs) by intraperitoneal injection. CFUs from the lungs, liver, and spleens on days 1, 28, and 56 post-infection were enumerated. A total of four mice per strain per timepoint were used in this experiment. Both lungs, the liver, and the spleen were homogenized for each sample at each timepoint. To homogenize the samples, organs were transferred to conical tubes that contained a 1:1 ratio of 1.6mm diameter and 0.9-2mm diameter stainless steel beads (Next Advance) and PBS with 0.05% Tween-80. Stainless steel UFO beads, 3.5mm (Next Advance) were also added for processing the lungs. Samples were homogenized in a Bullet Blender 5E Pro for 4 minutes at speed 10 setting. A 10-fold serial dilution was performed for each sample, and 100μl of each dilution was plated on 7H10 plates. Bacteria were grown at 37°C and colonies were counted two weeks later.

### Macrophage infection with *Mah* and *Mtb*

Primary murine macrophages were derived from the bone marrow of C57BL/6 mice by isolating cells from femurs. Bone marrow extracts were treated with 5ml ACK lysis buffer (Gibco) for 5 minutes and seeded onto non-tissue culture treated 15 cm plates at a density of 10^6^ cells per plate. Cells were plated in 20ml of DMEM supplemented with 10% FBS, 2mM glutamine (Gibco), 0.11 mg/ml sodium pyruvate (Gibco), and 10% MCSF derived from the supernatant of 3T3-MCSF cell culture. Macrophages were given an additional 10ml of media on days 3 and 6 and were differentiated 8 days total.

Differentiated macrophages were seeded at a cell density of 1.0 × 10^5^ cells per well in a 24-well plate 48 hours before infection for CFU enumeration and 0.5 × 10^6^ cells per well in a 6-well plate for RNA isolation. *Mah* and *Mtb* were grown to logarithmic phase, pelleted by centrifugation at 3500*g* for 5 minutes and washed with PBS twice. Cultures were then centrifuged at 200*g* to remove aggregates. Bacteria in single cell suspension were resuspended in macrophage media supplemented with 10% horse serum and allowed to opsonize for 10 minutes at room temperature. Macrophages were then ‘spinfected’ at a multiplicity of infection (MOI) of 1.5 for CFU enumeration or 5 for RNA isolation by centrifuging the cells and bacteria at 400*g* for 10 minutes. Cells were returned to macrophage media and daily half media changes were performed. For CFU enumeration, cells were washed once with PBS and lysed in water with 0.5% Triton-X. Following an incubation of 10 minutes, lysed cells were resuspended and serially diluted in 7H9 media. Dilutions were plated on 7H10 plates. For RNA isolation, cells were washed once with PBS, lysed in Trizol, and stored at −80°C until RNA extraction.

### Cytokine Bead Array Assay (CBA)

Supernatants from *Mah*-infected BMMs were harvested at 24 hours post-infection and filtered through 0.2 micron PVDF low-binding filters. Cytokine levels in supernatants were measured using the LEGENDplex mouse inflammation panel (Biolegend #740446) according to the manufacturer’s instructions with the following modifications. The recommended reagent and sample volumes were halved. Samples and standards were incubated with the beads overnight at 4°C. Samples and beads were stationary during incubations. Analyses were performed on a BD LSR Fortessa Analyzer.

### RT-qPCR

BMM infections for RT-qPCR were performed with an SmT-locked mc^2^2500 strain that contained multiple plasmid-encoded copies of *erp* and a matched SmO strain containing empty vectors. RNA was reverse-transcribed into cDNA with the SuperScriptIII-RT kit (Invitrogen) according to the manufacturer protocol, using 500ng input RNA per sample and oligo_d_T-primed amplification. A standard curve was generated using a separate reverse-transcribed sample composed of each individual sample mixed in equivalent amounts to total 500ng. This standard was diluted 1:6 in water for the top working standard, then diluted 5 times in 1:4 serial dilutions. Samples and NRT control were diluted 1:15 in water. Samples were assayed in biological and technical triplicate comparing the gene of interest to GAPDH transcripts on the same plate. The final reaction mix contained 10ul SsoAdvanced Universal SYBR Green Supermix (Biorad), 250nM of each primer, and 6ul of diluted cDNA template. Cycling was conducted with a 2-step amplification protocol (initial 95°Cx30 seconds, then 40 cycles of 95°Cx10 seconds and 60°Cx30 seconds, followed by a melt curve) in a CFX Connect thermal cycler (Biorad). Primer sequences were sourced from PrimerBank^101^ when possible.

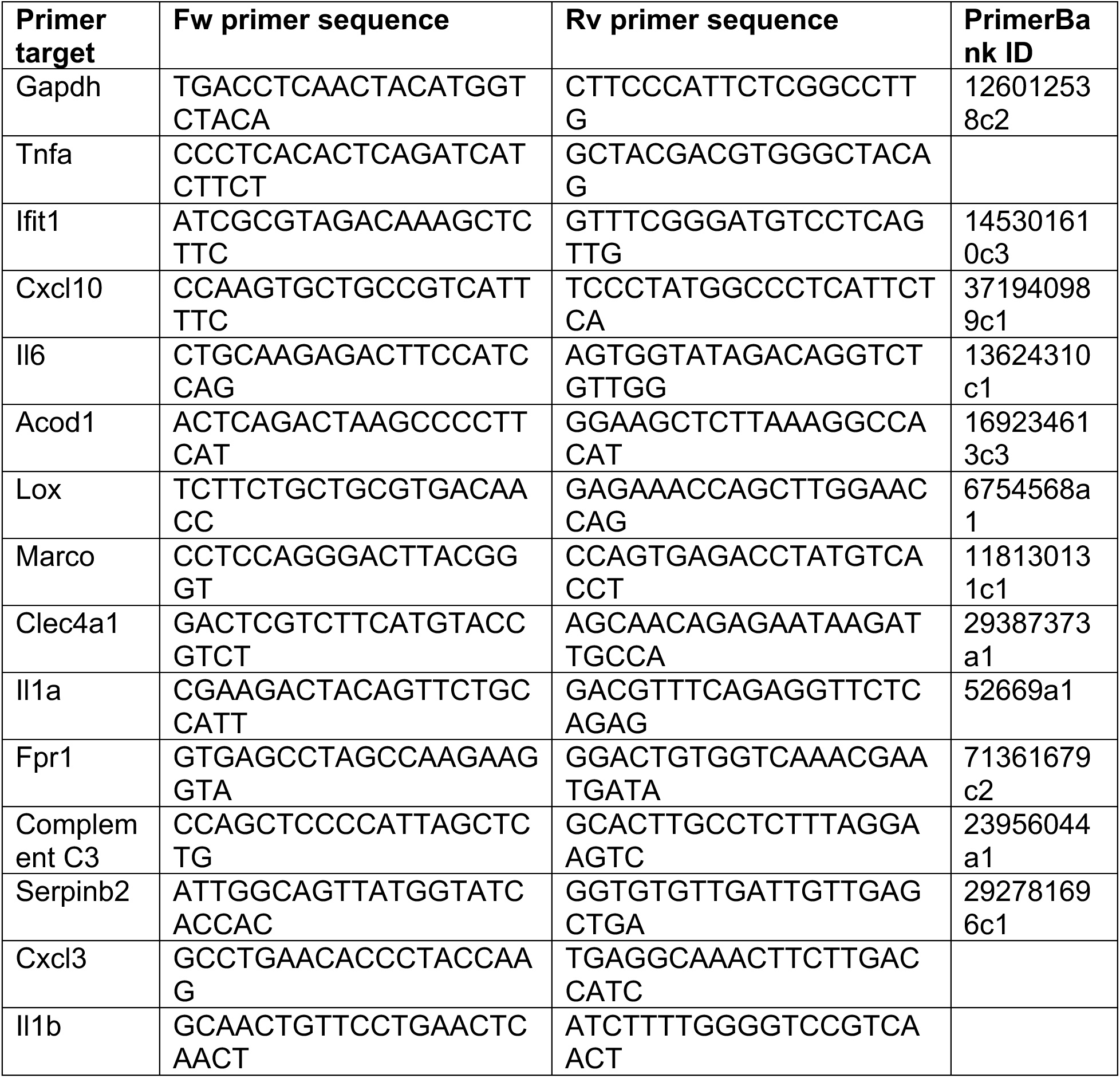

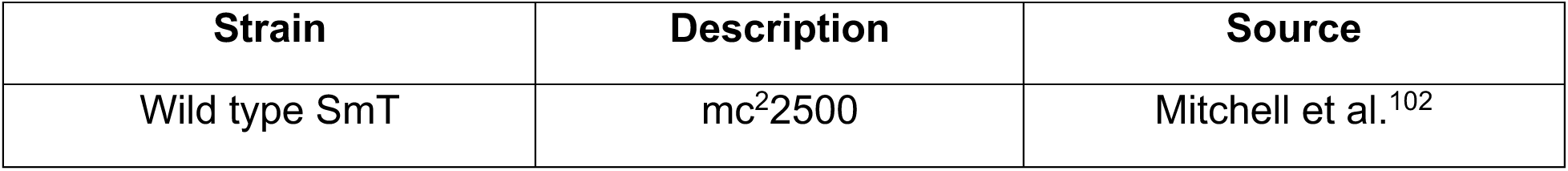
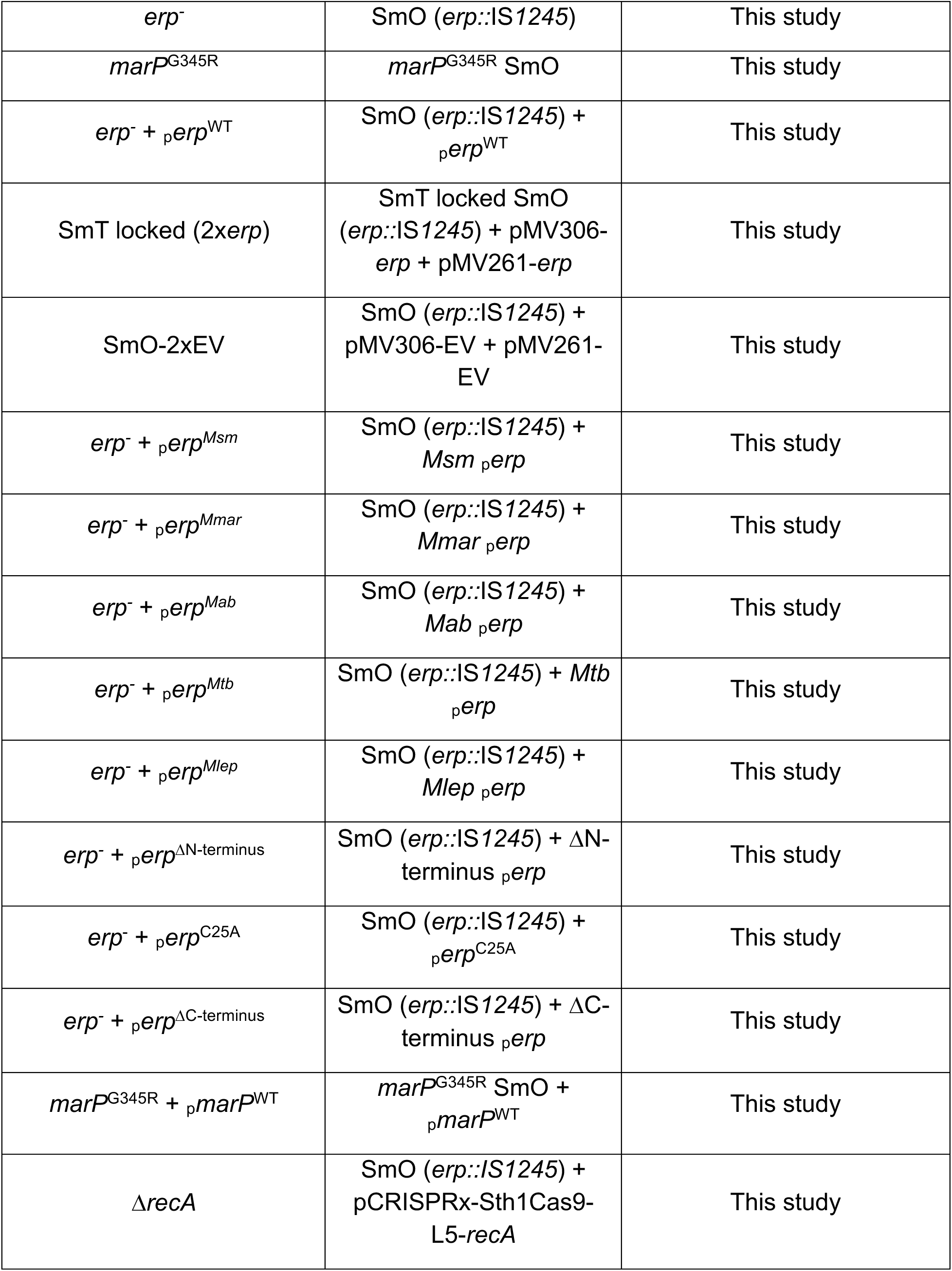
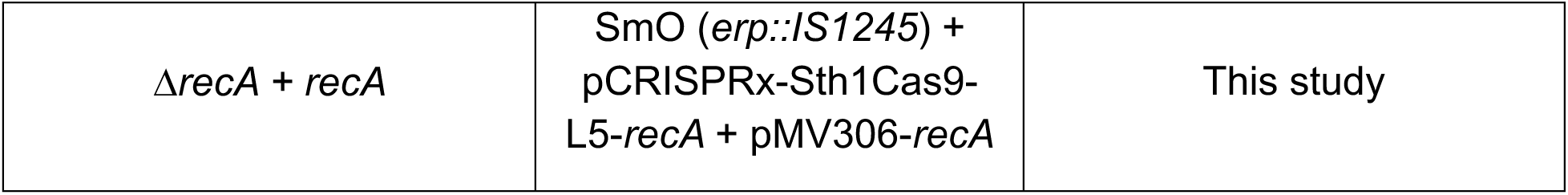

## Ethics statement

Mouse use and procedures were approved by the Office of Laboratory and Animal Care at the University of California, Berkeley (protocol AUP-2015-11-8096-1) in adherence with federal guidelines from the Guide for the Care and Use of Laboratory Animals of the National Institute of Health.

## Data availability

All DNA and RNA sequencing data generated in this work will be available through NCBI Sequence Read Archive (SRA) upon publication. *Mtb* RNA-seq data is available at https://artms.shinyapps.io/tb-omics-explorer/.

## Acknowledgements

We would like to thank the members of the Cox, Stanely, and Vance labs for critical feedback, with special thanks to Christopher Rae, Christopher Bennett, Varun Sridhar, and Anthony Rodriguez Vargas for experimental assistance and Christine Qabar for editorial feedback. We thank Azenta Life Sciences for RNA sequencing. This work was supported by National Institutes of Health grants U19AI162583 (J.S.C.) and U19AI135990 (J.S.C.), and the Henry Wheeler Center for Emerging and Neglected Diseases.

---

**Supplementary Figure 1.**
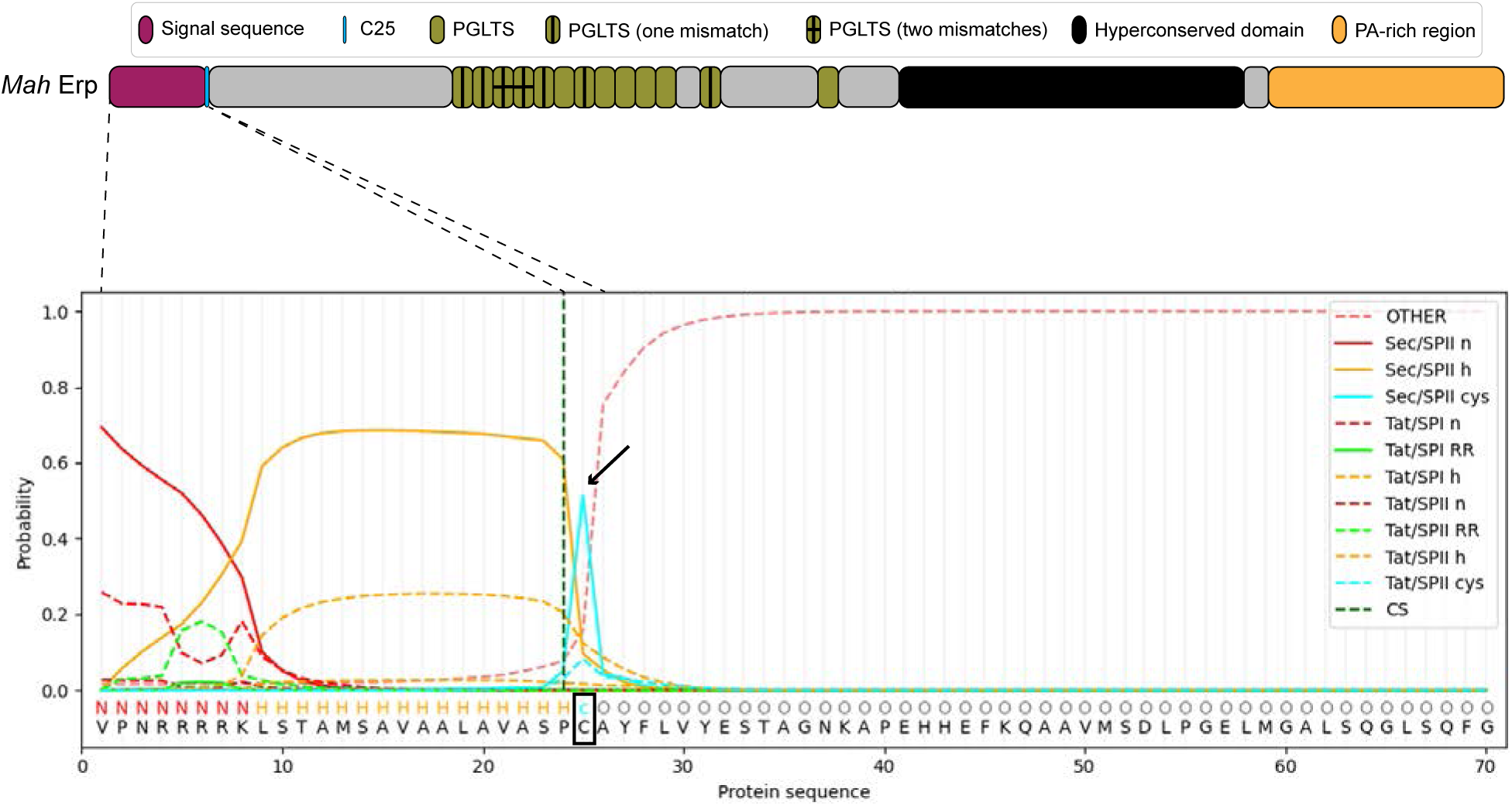
Predicted lipidation site of *Mah* Erp. SignalP 6.0 prediction of *Mah* Erp^49^. The cysteine residue (C25, boxed) is predicted to be a Sec substrate that is cleaved by SPase II as indicated by the light blue peak and arrow, which suggests that Erp is a prokaryotic lipoprotein. Dashed lines represent the protein feature that corresponds with the SignalP 6.0 prediction.

**Supplementary Figure 2.**
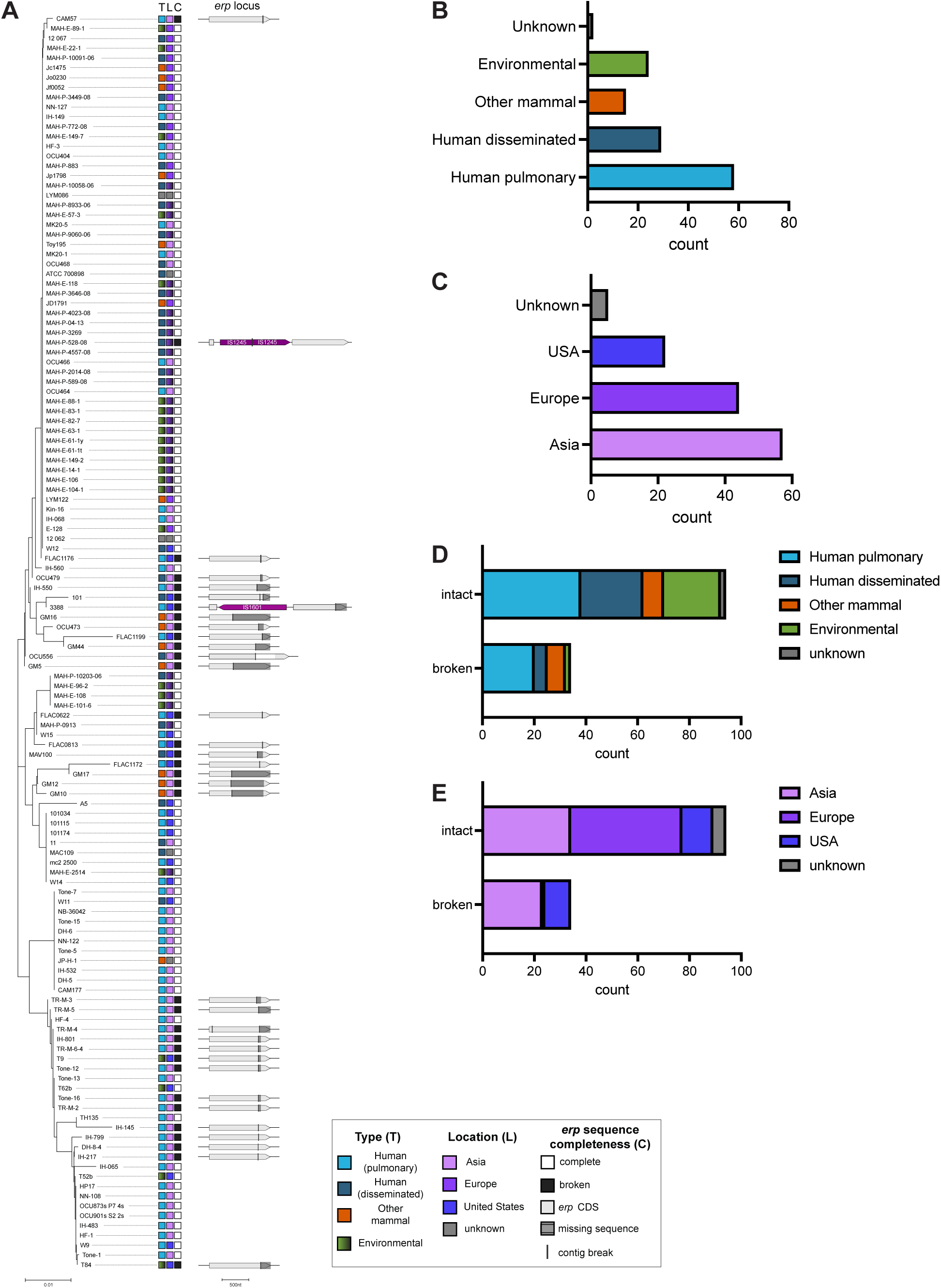
*Erp* sequence completeness in *Mah* isolate genome sequences. **A.** Minimum evolution phylogeny of *erp* coding sequences (DNA) for all *Mah* genome sequences in Genbank. Metadata regarding *Mah* isolate source and location is indicated by squares in the first and second column. The third column indicates whether the *erp* sequence was found to be continuous on one contig (complete) or split across two contigs (broken). If the *erp* sequence was broken or incomplete, the gene diagrams to the right indicate the position of the contig break, and missing sequence is represented by grey translucent rectangles. **B.** Number of *Mah* isolate genomes from each isolate source. **C.** Number of *Mah* isolate genomes from each isolation location. **D.** and **E.** *Mah* isolate counts by source and location that contained broken versus intact *erp* sequences.

**Supplementary Figure 3.**
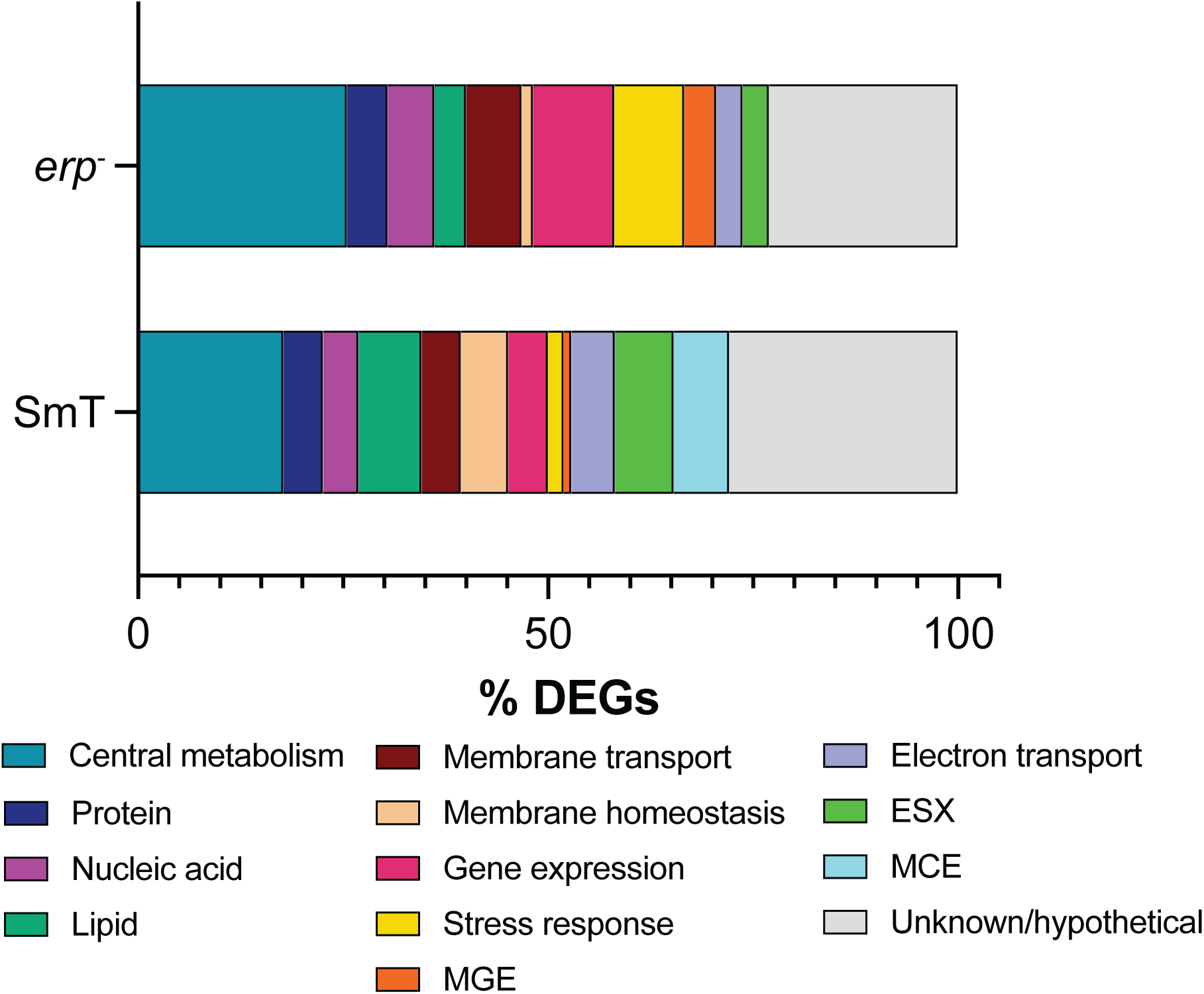
Gene ontology analysis of differentially expressed genes in the SmT and *erp*^−^ morphology. Stacked bar graph representing the proportion of differentially expressed genes (DEGs) representing each considered gene ontology category. DEGs from each individual comparison are pooled in this analysis. Categories represent gene ontology metabolic processes unless otherwise denoted. MGE: mobile genetic element. ESX: type VII secretion system. MCE: mammalian cell entry.

**Supplementary Figure 4.**
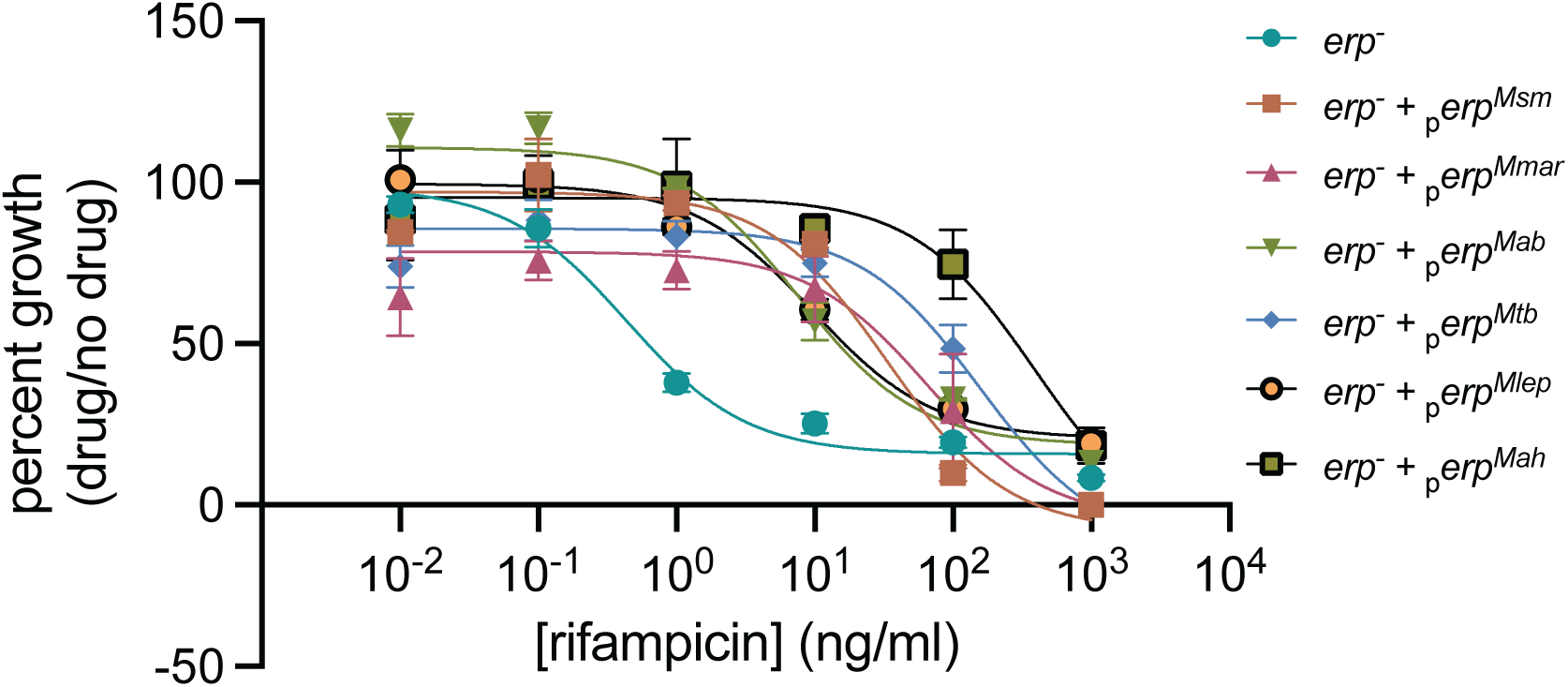
Different mycobacterial *erps* restore *erp*^−^ resistance to antibiotics to varying degrees. Dose response curves for *erp*^−^ and *erp*^−^ + _p_*erp*^Msm^, _p_*erp*^Mmar^, _p_*erp*^Mab^, _p_*erp*^Mtb^, _p_*erp*^Mlep^, and _p_*erp*^Mah^ against rifampicin measured as OD_600_ of the culture grown with the rifampicin at the indicated concentration divided by OD_600_ of the culture grown without rifampicin. Each data point is the mean ± standard deviation of two technical replicates. The curve is representative of two biological replicates.

**Supplementary Figure 5.**
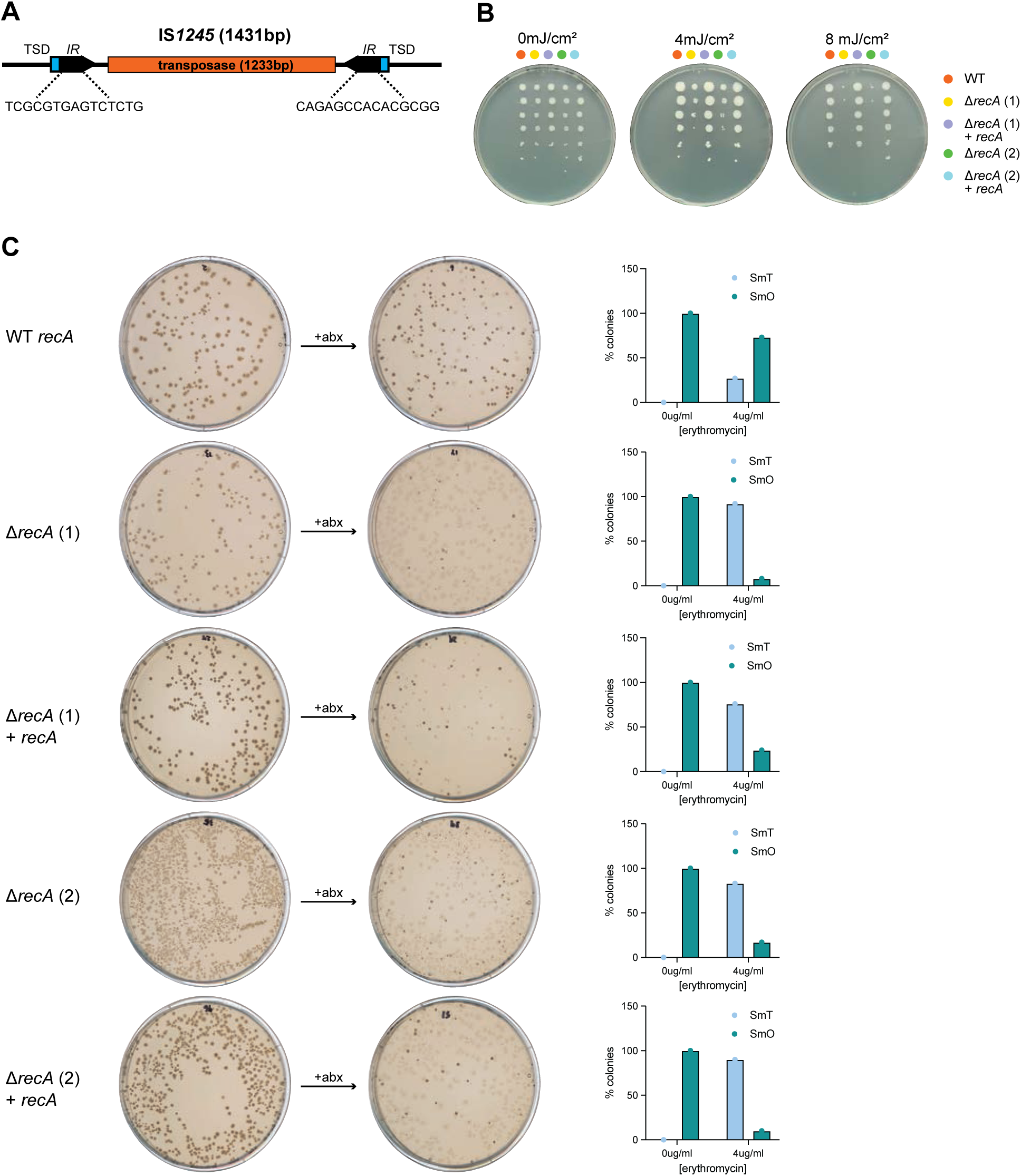
IS*1245* transposition is RecA-independent. **A.** Schematic of IS*1245*. Target site duplications (TSD) and inverted repeats (IR) are indicated on the diagram. **B.** UV exposure assay of WT *Mah*, Δ*recA* (independent clones 1 and 2), and Δ*recA* + *recA*. Amount of UV bacteria were exposed to is indicated above each plate. **C.** Switching assay for WT *Mah*, Δ*recA*, and Δ*recA* + *recA*. Representative plate scans of WT *Mah*, Δ*recA*, and Δ*recA* + *recA* treated with 0μg/ml or 4μg/ml of erythromycin. The percentage of colonies with SmO and SmT morphotypes after treatment with 0 μg/ml or 4μg/ml of erythromycin is quantified to the left of each plate.

**Supplementary Figure 6.**
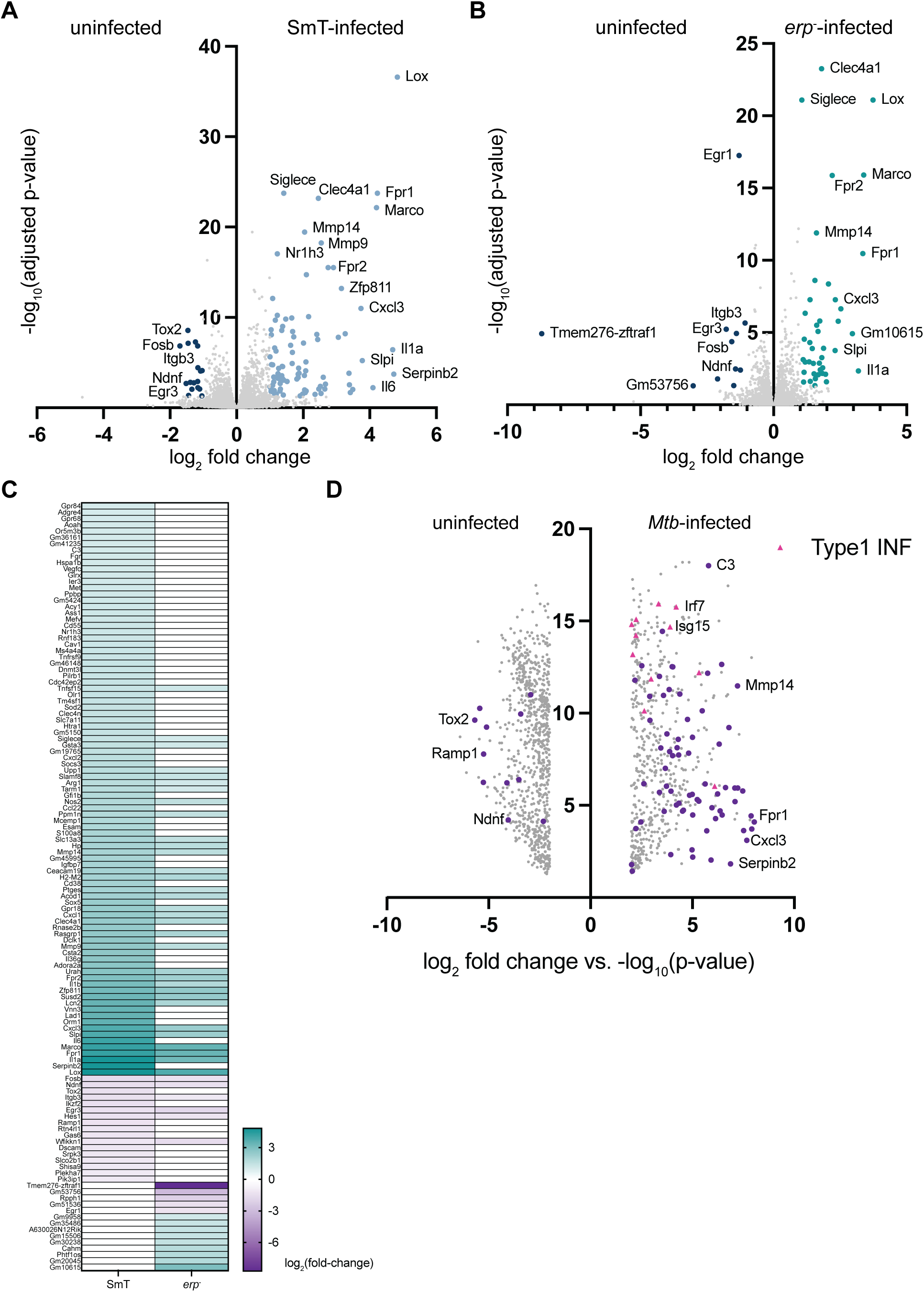
Characterization of the immune response of BMMs infected with *Mah*. **A.** Volcano plot of differentially expressed genes in SmT-infected BMMs compared to uninfected controls. Adjusted p-value ≤ 0.05 and log_2_-fold change ≥ 1.0 were considered significant and represented in light blue (upregulated) and navy (downregulated), nonsignificant results represented in grey. Select genes of interest are labeled on the plot. **B.** Volcano plot of differentially expressed genes in *erp*^−^-infected BMMs compared to uninfected controls. Adjusted p-value ≤ 0.05 and log_2_-fold change ≥ 1.0 were considered significant and represented in teal (upregulated) and navy (down-regulated), nonsignificant results represented in gray. Select genes of interest are labeled on the plot. **C.** Heat map of the relative gene expression of all the differentially expressed genes in SmT- and *erp*^−^-infected BMMs. **D.** Volcano plot of differentially expressed genes in *Mtb*-infected BMMs compared to uninfected controls at 24 hours post-infection. Adjusted p-value ≤ 0.05 and log_2_-fold change ≥ 2.0 were considered significant and represented on the plot. Differentially expressed genes shared with *Mah*-infected BMMs are highlighted in purple and type 1 interferon (IFN) genes are highlighted in pink.

**Supplementary Figure 7.**
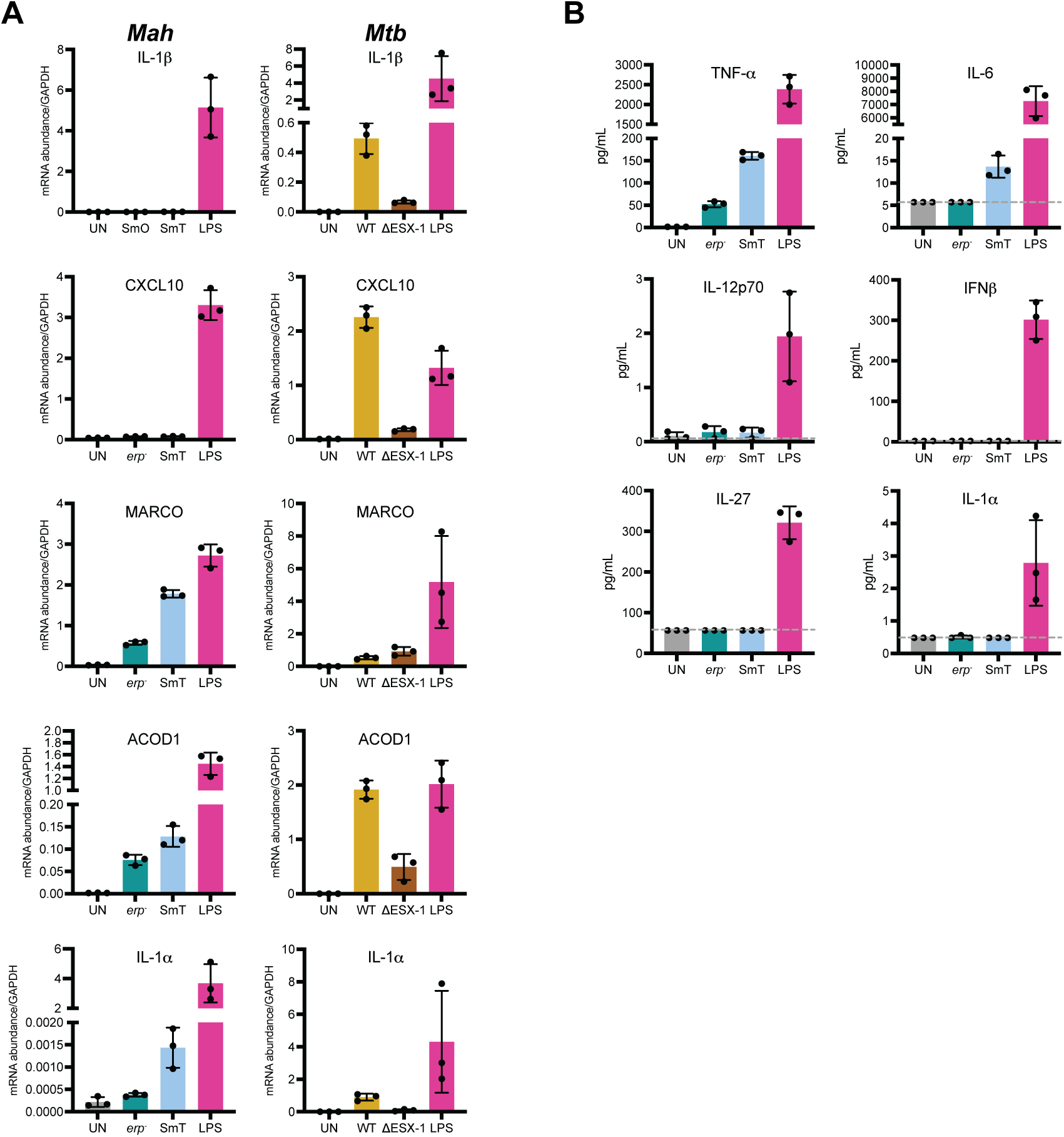
The immune response of *Mah*- and *Mtb*-infected BMMs differ. **A.** RT-qPCR of select transcripts in SmT *Mah*, *erp*^−^ *Mah*, WT *Mtb*, and ΔESX-1 *Mtb*-infected macrophages 24 hours post-infection. mRNA levels are normalized to GAPDH. Each data point is the mean ± standard deviation of three biological replicates. **B.** Cytokine levels in the supernatants of SmT *Mah* and *erp*^−^ *Mah*-infected macrophages 24 hours post-infection quantified using the LEGENDplex mouse inflammation panel for cytokine bead array assays. Each data point is the mean ± standard deviation of three biological replicates. Gray dashed lines indicate lower limits of detection.

**Supplementary Figure 8.**
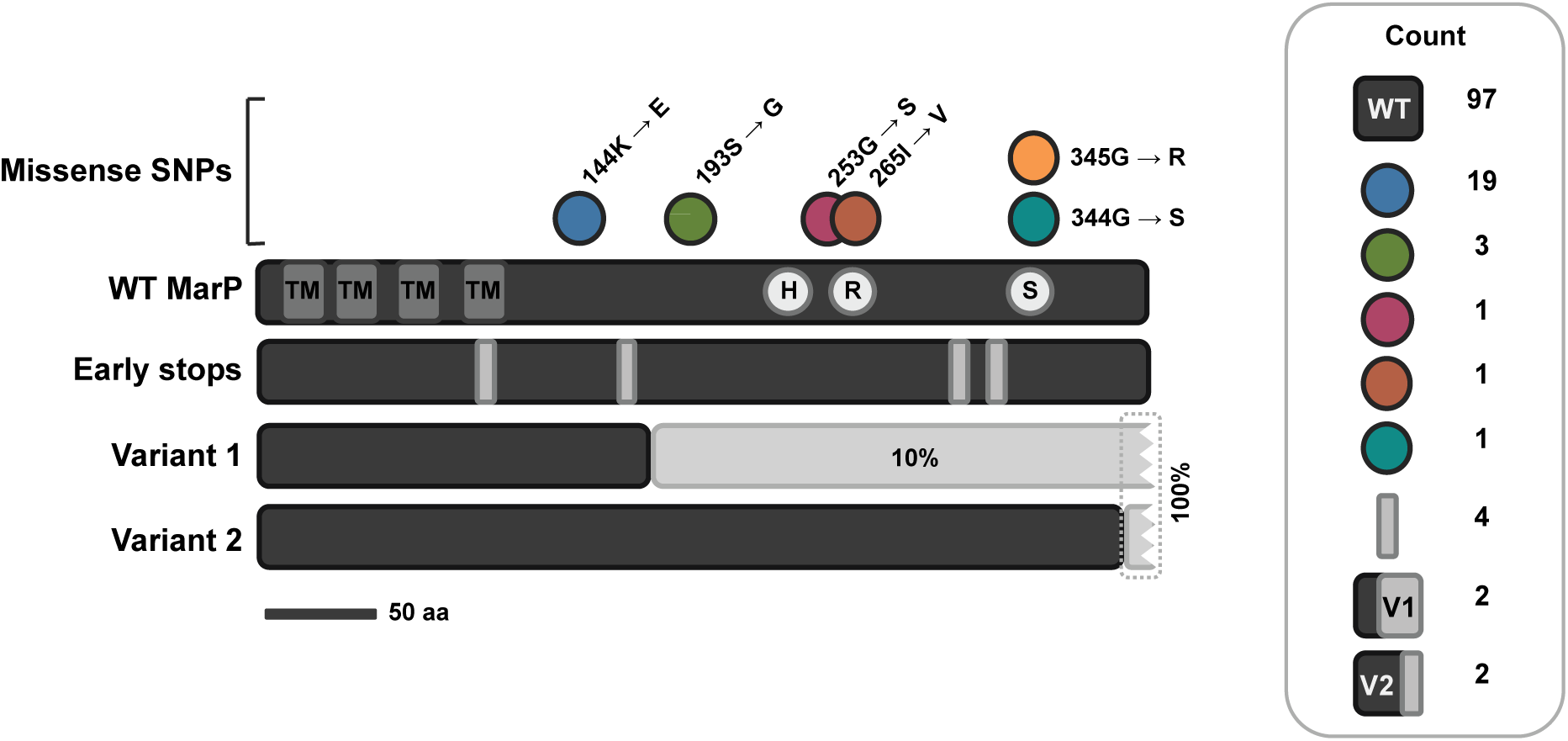
Variation in MarP protein sequence among *Mah* isolates. Protein domain architecture of wild-type MarP sequence is indicated on the top black bar (TM: transmembrane domain, H, R, and S: active site residues). Dots above indicate the position and nature of various nonsynonymous SNPs detected in some *Mah* isolate sequences (strains with these variations contained only one of the listed SNPs). The yellow dot represents the MarP SNP derived from mc^2^2500 in this work. MarP sequences with nonsense mutations resulting in early stop codons are indicated on the early stops bar, each grey bar indicates the position of a different strain’s mutation. Bars labeled variants 1 and 2 represent alleles of MarP that have substantial sequence variation induced by deletions that shift the reading frame but do not introduce early stop codons. These variant sequences were found on ends of contigs without stop codons, represented by the jagged break at the end of the bar. Rectangle labeled “count” shows the number of *Mah* isolates encoding each MarP allele detected in this analysis.

**Supplementary Figure 9.**
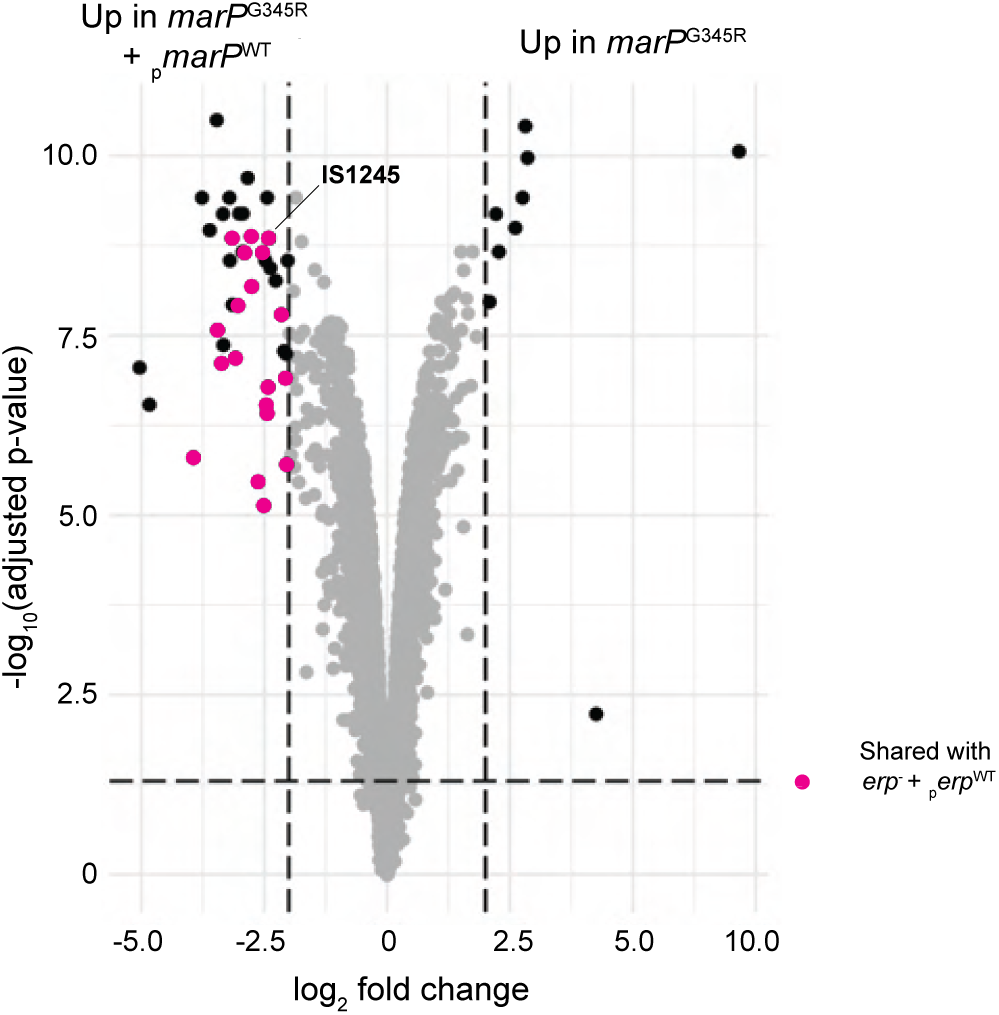
The transcriptional profile of *marP*^G345R^ and *marP*^G345R^ + _p_*marP*^WT^. Volcano plot of differentially expressed genes in *marP*^G345R^ + _p_*marP*^WT^ compared to *marP*^G345R^. Adjusted p-value ≤ 0.05 and log_2_-fold change ≥ 2.0 were considered significant and represented in black and nonsignificant results represented in grey. Differentially expressed genes that are shared between *marP*^G345R^ + and *erp*^−^ + _p_*erp*^WT^ are highlighted in pink. _p_*marP*^WT^

